# YY1 protein is essential for the promotion of Muller glia reprogramming and retina regeneration

**DOI:** 10.1101/2025.01.31.635905

**Authors:** Mansi Chaudhary, Omkar Mahadeo Desai, Poonam Sharma, Sharanya Premraj, Pooja Shukla, Rajesh Ramachandran

## Abstract

Unlike mammals, the Muller glia reprogramming in zebrafish retina restores vision after an acute retinal injury. Here, we explored the Ying yang (Yy1) protein and its multi-faceted roles in different phases of retina regeneration. We show that the acetylation and deacetylation status of Yy1 contribute to its transcriptional activation and repression functions on various target genes, including regeneration-associated genes (RAGs). Yy1 is regulated positively by TGF-β and negatively through Delta-Notch signaling in the injured retina. Yy1-knockdown caused reduced retinal progenitor induction and regeneration, while the opposite was seen in its overexpression. Yy1 collaborates with histone deacetylases, BAF complex, and the effector of TGF-β signaling, pSMAD3, to target the genome differentially. Lastly, the whole transcriptome analysis of the Yy1-debilitated retina revealed differential expression of various RAGs and BMP-signaling. Yy1 facilitates the BMP pathways genes through the downregulation of *noggin3*. Our study unravels how a single transcription factor, Yy1, could influence many important regulatory steps of retina regeneration.

## Introduction

Vertebrates such as zebrafish possess remarkable regenerative capacity of various organs after an acute injury, including parts of the central nervous system^1, 2^. The retina is part of the central nervous system, which regenerates in zebrafish irrespective of the mode of injury^3^. Unraveling the mechanisms of zebrafish retina regeneration has been pivotal in emulating regenerative response in mammalian systems. In general, mammalian models have resorted to wound healing over regeneration, as observed in amphibians and fish^4^. While various regenerative pathways unraveled in zebrafish have resulted in similar positive responses in mammalian models^5–9^, the enigma of regeneration remains far from clear. Despite inadequacies in completing the regenerative responses, the regeneration mechanism draws parallels among vertebrates ^3, 10, 11^. Upon injury, zebrafish retinal Müller glia undergoes reprogramming through hierarchical gene regulatory networks, including the induction of pluripotency-inducing factors^12–19^ resulting in the formation of retinal progenitors. The *de novo*-induced retinal progenitors expand in number and differentiate back to various retinal neurons and Müller glia^20^. During this regeneration process, various genetic￼ and epigenetic factors￼ come into action ^20, 21^. The factors that contribute to progenitor induction often include tumor suppressors such as PTEN^12^, epigenome modifiers such as Hdacs^16^, post-transcriptional gene regulator miRNAs^13, 14, 17, 19, 22^

We explored the importance of the zinc finger transcription factor Ying yang 1 (Yy1), ubiquitously expressed in all tissues^23^. Yy1 is known for its dual role as a transcriptional activator and repressor on different target genes^24^. Yy1 is a highly conserved gene from Drosophila, *C*. *elegans,* zebrafish, and mouse to humans. Deletion of Yy1 is embryonically lethal in mice ^25^. Yy1 protein is stable in its phosphorylated form in all cell types ^23^ suggesting the bifunctional nature of Yy1 is probably attributed to its post-translational modifications ^26–28^ and collaborating partners^29–31^. Yy1 is closely associated with epithelial to mesenchymal transition and maintaining stemness in cancer cells ^32–34^. Yy1, along with CTCF, enhances axon regeneration^35^ and X-chromosome inactivation^36^. YY1 governs the multidimensional epigenetic landscape of embryonic stem cells through its ability to modulate DNA methylations and histone modifications^37^. Yy1 is involved in the heart^38^, muscle^39, 40^, axon^35^, and liver^41^ regeneration in various model systems. However, little is known about the roles of Yy1 during the zebrafish retina regeneration.

Here, we explored the expression dynamics of *yy1a* and *yy1b* in zebrafish retina and its significance during regeneration through knockdown and overexpression. We saw the differential influence of Yy1 on the expression of regeneration-associated genes (RAG) in the damaged zebrafish retina. We also unraveled the functional importance of Yy1 in conjunction with the BAF complex, an ATP-dependent chromatin remodeler implicated in development and diseases^42^. Whole transcriptome analysis of yy1-knocked down regenerating retina revealed its importance in governing different cell signaling and immune system pathways. Lastly, we demonstrated the unique roles of Yy1 in its acetylated and deacetylated status during retina regeneration. Together we could prove that Yy1-derived gene regulatory and cell signaling pathways are inevitable for normal retina regeneration in zebrafish. Previous studies have demonstrated the importance of Tgf-β signaling during retina regeneration. In this study, we show that the *yy1* gene is regulated via Tgf-β signaling, which in turn is essential to maintain the BMP signaling during retina regeneration.

## Results

### Yy1 is downregulated in post-injured retina

We explored the levels of zebrafish Ying yang paralogs *yy1a* and *yy1b* at mRNA and protein levels at various times post-retinal injury. Yy1a and Yy1b show approximately 88% protein similarity (Figure S1A). We saw a sudden decline in the mRNA (Figure 1A) and protein (Figure 1B) levels of *yy1a* and *yy1b* from 12hpi to 2dpi. As the cellular proliferation started at 36hpi, the *yy1* levels started showing a steady upward trend, reaching up to uninjured control levels by 4dpi, when proliferation is at its peak in the needle poke injury method. This observation suggested that Yy1 levels get downregulated, specifically during the reprogramming phase of the retina regeneration. We further explored the *yy1a* (Figures 1C and S1B) and *yy1b* (Figures 1E and S1C) mRNA levels at various times post-retinal injury to ascertain the correlation of their expression with regard to cellular proliferation. Three selected time points of 2dpi, 4dpi, and 8dpi were used to analyze the expression of *yy1a* and *yy1b* as these points marked the return of *yy1* RNA from the downregulated state seen at 12hpi to 2dpi time window. The co-labeling of BrdU and *yy1a* (Figure 1D) and *yy1b* (Figure 1F) reveals that most proliferating cells do not express *yy1a* or *yy1b*. Of those proliferating cells with *yy1* expression, the *yy1a* correlates more with BrdU^+^ cells than *yy1b* at earlier time points. We decided to explore further the expression pattern of *yy1a* and *yy1b* by making use of the transgenic line *1016tuba1a*:GFP, which, upon injury, selectively marks the Müller glia-derived progenitors with GFP. Using a fluorescent-based cell sorting approach, we used this 4dpi transgenic retina to sort the GFP+ and GFP-cells. The isolated cells are used to analyze expression levels of *yy1a* and *yy1b* RNAs. The qPCR analysis of *yy1a* and *yy1b* levels reveals that the Müller glia cells from the regenerating retina have reduced levels of *yy1a* and *yy1b* at 4dpi (Figure 1G). Analysis of Yy1 protein on 4dpi and 6dpi retinal sections also showed a reduction in its expression in BrdU^+^ proliferating cells (Figures 1H, 1I, and S1D). Based on these results, it is safe to assume that the return of *yy1a* and *yy1b* during the proliferative phases of retina regeneration in cells other than the BrdU^+^ cells.

**Figure 1.**
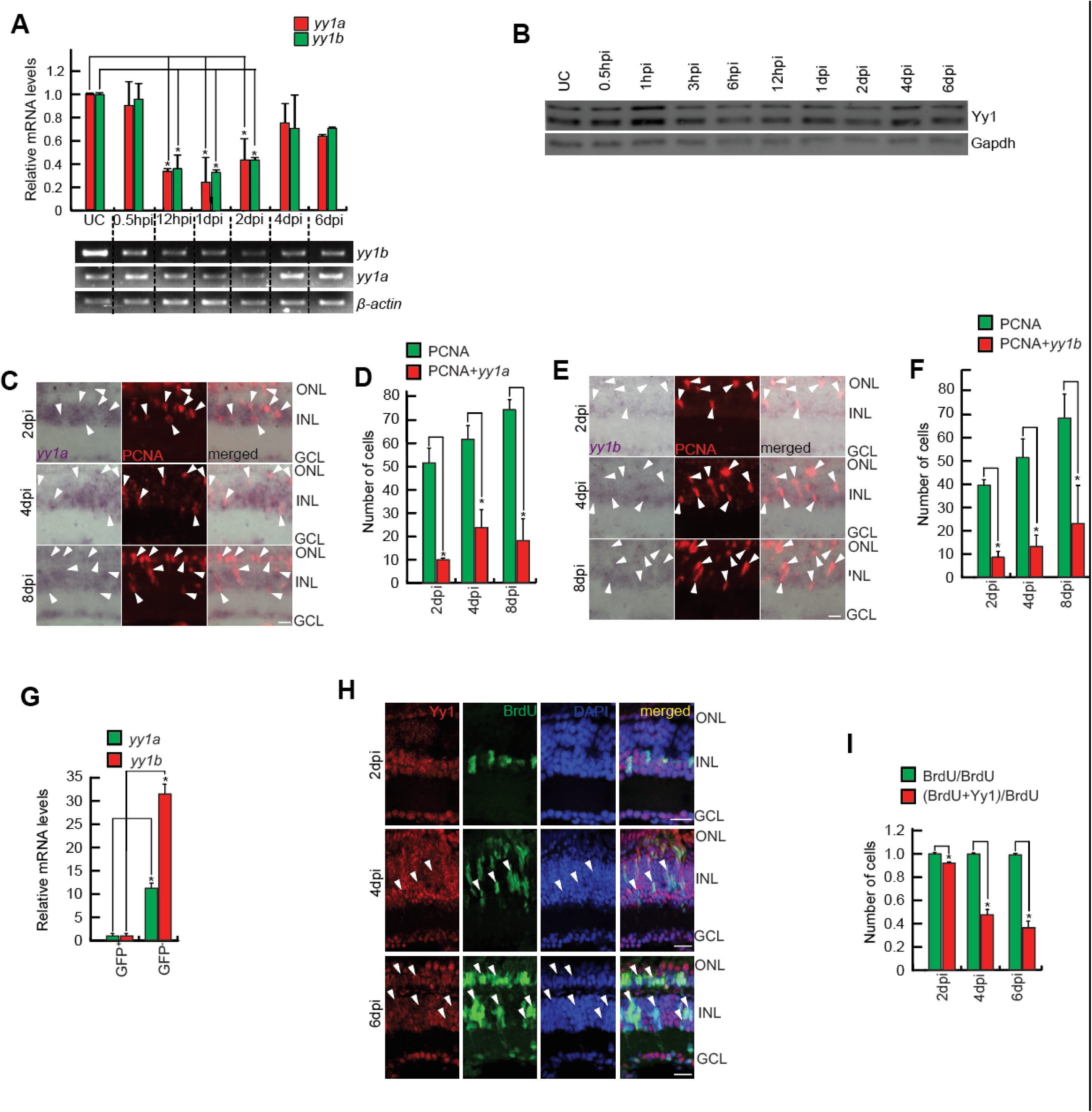
Yy1 is ubiquitously expressed in the entire retina and absent from majority of proliferating retinal progenitors. (**A**). The qPCR (upper) and RT-PCR (lower) analyses of *yy1a and yy1b* genes in the retina at various time points post-retinal injury; *p < 0.0001; n=6 biological replicates. **(B)** Western Blot analysis of Yy1 from retinal lysates prepared at different time points post-injury. Gapdh is the loading control. **(C and D)** The 60X bright field (BF) and fluorescent images of retinal cross-sections show the exclusion of *yy1a* mRNA from the proliferating cells which are marked by PCNA at 2dpi, 4dpi and 8dpi (C), which is quantified in (D); *p < 0.005; n=6 biological replicates. **(E and F)** The 60X bright field (BF) images of retinal cross-sections show the exclusion of *yy1b* mRNA from the proliferating cells which are marked by PCNA at 2dpi, 4dpi and 8dpi (E), which is quantified in (F); *p < 0.005; n=6 biological replicates. White arrow heads in **C**, and **E** marks PCNA positive but *yy1a*/*yy1b* negative cells. **(G)** The qPCR analysis shows high levels of *yy1a* and *yy1b* in GFP^-^ as compared to GFP^+^ fraction of cells from *1016 tuba1a: GFP* *p<0.0005; n=6 biological replicates. **(H and I)** The 60X Immunofluorescence (IF) microscopy images of retinal cross sections show exclusion of Yy1 from the proliferating cells marked by BrdU at 2dpi, 4dpi and 6dpi (H), which is quantified also (I) *p < 0.0001; n=6 biological replicates. Scale bars represent 10μm in (C, E and H); asterisk marks the injury site and GCL, ganglion cell layer; INL, inner nuclear layer; ONL, outer nuclear layer in (C, E and H); dpi, days post injury.

### Yy1 is essential for normal retina regeneration

We decided to explore whether Yy1 is important for the retina regeneration. In the injured retina, we performed a morpholino-based gene knockdown of the *yy1a* and *yy1b* genes (Figure 2A). The retinal sections after *yy1a* and *yy1b* gene knockdowns using antisense morpholinos (MOs) alone and in combination caused a reduction in the proliferating population of BrdU^+^ or PCNA^+^ retinal progenitors at 4dpi compared to control conditions

**Figure 2.**
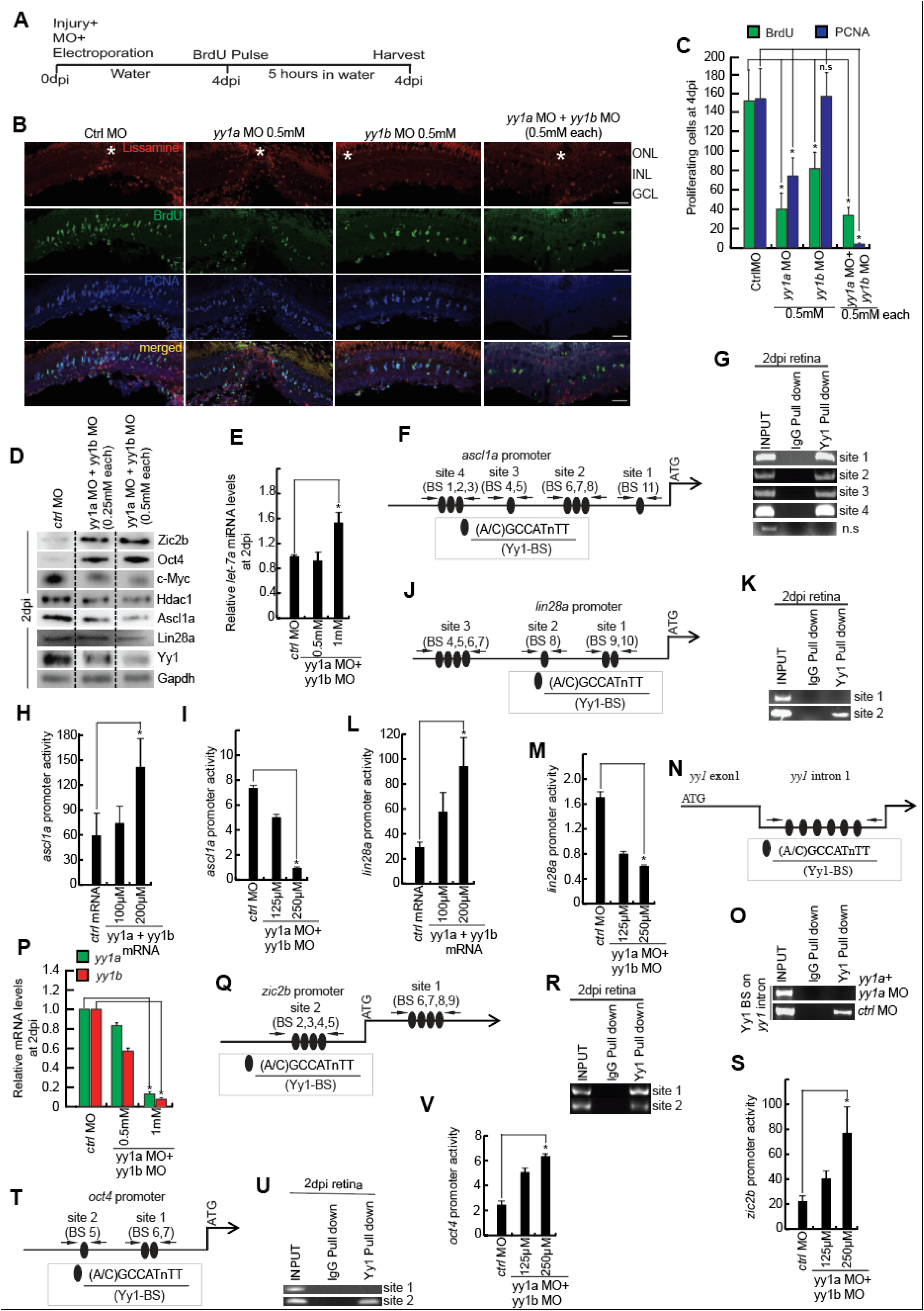
Yy1 has a pro-proliferative role post-retinal injury and regulates the expression of several regeneration-associated genes (RAGs). **(A)** An experimental timeline that describes injury, Lissamine-tagged Morpholino (MO) delivery, electroporation at 0dpi, BrdU pulse for 5hrs at 4dpi, followed by harvesting. **(B and C)** IF microscopy images of retinal cross-sections show a decrease in the number of BrdU^+^ retinal progenitors with *yy1a* MO and *yy1b* MO alone and in combination, compared to control MO injected retina at 4dpi (B), which is quantified (C). **(D)** Western Blot analyses of various RAGs in retinal extracts prepared from retinae electroporated with MOs against *yy1a* and *yy1b* at 2dpi. (E) The qPCR analyses of *let-7a* mi-RNA in the *yy1a* and *yy1b* knockdown condition at 2dpi; *p < 0.05; n=6 biological replicates. **(F and G)** The schematic showing Yy1-binding sites (BS) on *ascl1a* promoter (F) and the retinal ChIP assays confirm the physical binding of Yy1 on *ascl1a* promoter, in 2dpi retina (G). N.S marks the negative control. **(H and I)** Luciferase assay done in zebrafish embryos at 24hpf, injected with *ascl1a: GFP-luciferase* construct shows upregulation in the *ascl1a* promoter activity in *yy1a* and *yy1b* overexpression (H), while downregulation in their combined knockdown (I). (**J and K)** The schematic showing Yy1-binding sites (BS) on *lin28a* promoter (J) and the retinal ChIP assays confirm the physical binding of Yy1 on *lin28a* promoter, in 2dpi retina (K). **(L and M)** Luciferase assay done in zebrafish embryos at 24hpf, injected with *lin28a: GFP-luciferase* construct shows upregulation in the *lin28a* promoter activity in *yy1a* and *yy1b* overexpression (L), while opposite in their combined knockdown (M). **(N and O)** The schematic showing Yy1-binding sites (BS) on *yy1a* intron (N) and the retinal ChIP assays confirm the loss of Yy1 binding onto its intron, in 2dpi retina (O). **(P)** The qPCR analysis shows the downregulation of *yy1a* and *yy1b* in their own combined knockdown. **(Q and R)** The schematic showing Yy1-binding sites (BS) on *zic2b* promoter (Q) and the retinal ChIP assays confirm the physical binding of Yy1 on *zic2b* promoter, in 2dpi retina (R). **(S)** Luciferase assay done in zebrafish embryos at 24hpf, injected with *zic2b: GFP-luciferase* construct shows upregulation in the *zic2b* promoter activity in *yy1a* and *yy1b* knockdown. **(T and U)** The schematic showing Yy1-binding sites (BS) on *oct4* promoter (T) and the retinal ChIP assays confirm the physical binding of Yy1 on *oct4* promoter, in 2dpi retina (U). **(V)** Luciferase assay done in zebrafish embryos at 24hpf, injected with *oct4: GFP-luciferase* construct shows upregulation in the *oct4* promoter activity in *yy1a* and *yy1b* knockdown. Error bars represent SD. *p < 0.001 in (H, L, M, P, S, V); *p < 0.05 (F). n = 6 biological replicates (C, E and P).

(Figures 2B and 2C). It is interesting to note that the knockdown of *yy1a* had a profound negative effect on negatively regulating cell proliferation than that of *yy1b*, while the combined *yy1a* and *yy1b* knockdown had an additive effect (Figure 2C). We then explored if the reduced progenitor proliferation in *yy1* knockdown conditions influenced regeneration-associated genes (RAG) such as Ascl1a￼, c-Myc￼, Hdac1￼, Oct4￼, and Zic2b^17^ (Figure 2D). We saw a decline in the levels of Ascl1a, c-Myc, Hdac1, and Yy1 itself but an unanticipated increase in the Zic2b and Oct4. One of the microRNAs, *let-7*, whose downregulation is essential for mRNA translation of several RAGs^19^ and Shh-signaling components, ￼ is upregulated in the *yy1* knockdown retina (Figure 2E). These elevated let-7 levels probably could account for reduced protein abundance of Ascl1a and c-Myc, which are post-transcriptionally silenced by this microRNA^19^. We saw an early decline and subsequent restoration of *yy1a* and *yy1b* levels in the regenerating retina. Based on this observation, we assessed the effect of a late knockdown of *yy1*. For this experiment, *yy1a* and *yy1b* MO were delivered at the time of injury but electroporated only on 4dpi after pre-labeling the progenitors with BrdU (Figure S2A). The progenitors were then labeled with a second marker EdU before retina harvest at 9dpi and immunofluorescence of the tissue sections (Figure S2B). The goal of this experiment was to address if the reappearance of *yy1* in the later phase of regeneration had any role in the progenitor amplification and differentiation. The results from BrdU^+^/EdU^+^ progenitor cell counting showed a reduced number of BrdU^+^/EdU^+^ double-positive cells, indicating that *yy1a* and *yy1b* combined knockdown reduced the continued proliferation of progenitors (Figure S2C). These results suggested that despite *yy1* being secluded from actively proliferating cells, their expression in the neighborhood may be important in driving progenitor proliferation.

We adopted a pharmacological approach to inhibit Yy1 function through nitric oxide (NO) donor DETA/NONOate. The S-Nitrosation of the Yy1 protein negatively affects its DNA-binding ability and normal function^43^. We used two different concentrations of DETA/NONOate to dip the retinal injured zebrafish (Figure S2D). The progenitor cell number declined in a DETA/NONOate in a concentration-dependent manner in the 4dpi retina (Figures S2E and S2F). We explored the levels of phospho-histone 3 (PH3), a marker of cellular mitosis. We saw a decline in the PH3^+^ cells in the DETA/NONOate-treated retina (Figure S2G and S2H). The DETA/NONOate treatment did not cause increased cell death, as seen by the TUNEL assay (Figure S2I). As the Yy1 protein is known for its transcriptional activation and repression roles, we explored if the functional blockade of Yy1 had any roles in the expression levels of various RAGs. Of the RAGs tested, we saw a moderate increase in the protein levels of Oct4, Zic2b, and Lin28a, while a decline in Ascl1a (Figure S2J). The levels of cMyc and Hdac1 remained unaffected. This result suggested that functional blockade of Yy1 has differential effects on the RAGs with a negative correlation with retinal progenitor proliferation.

### Yy1 regulates the expression of regeneration-associated genes

Previous studies have demonstrated the importance of several transcription factor genes essential for retina regeneration. Analysing a few regulatory DNA sequences, such as *ascl1a*, *lin28a*, *zic2b* and *oct4*, revealed consensus Yy1 binding DNA on their promoter sequences (Figure 2F, 2J, 2Q and 4T). We explored if Yy1 had any regulatory role in these gene expressions. Since Yy1 can influence gene expression positively or negatively, we first explored the levels of these genes after *yy1* knockdown (Figure S2K). We analyzed the promoters of these genes for potential Yy1 binding at its putative binding sites. A chromatin immunoprecipitation (ChIP) assay was adopted on 2dpi retinal tissue to confirm the Yy1 binding on the DNA regulatory sequences. On the *ascl1a* promoter, we saw eight potential Yy1 binding sites in four groups (Figure 2F), which bound to Yy1 as observed in the ChIP assay (Figure 2G). To ascertain this further, we performed luciferase assay in zebrafish embryos co-injected with *ascl1a* promoter driving EGFP-luciferase fusion construct and *yy1a* and *yy1b* mRNA or *yy1a* and *yy1b* morpholinos (MOs) in separate experiments. We saw a *yy1* mRNA concentration-dependent upregulation of *ascl1a* promoter activity (Figure 2H) and an anticipated downregulation with increasing concentrations of *yy1a* and *yy1b* MOs (Figure 2I). We saw a similar trend with the lin28a promoter, albeit a reduced number of Yy1 binding sites on its promoter (Figures 2J-2M). These results supported the direct involvement of Yy1 in positively regulating both *ascl1a* and *lin28a* in zebrafish. Similarly, Yy1a played the role of a positive regulator of its own gene expression. The Yy1 is bound to its binding sites present on the gene promoter, which is confirmed by ChIP and qPCR assays (Figures 2N-2P). Contrary to this, the Yy1 seemed to play a repressive role on the *zic2b* and *oct4* gene expression. The putative binding sites of *zic2b* promoter were functional in the ChIP assay with a concordant promoter activity revealed by luciferase assay in zebrafish embryos co-injected with *yy1a* and *yy1b* MO along with *zic2b* promoter driving luciferase (Figure 2Q-2S). The second Yy1-binding sites on *oct4* promoter was functional while the one closer to ATG start codon did not bind with Yy1 as observed in a 2dpi retinal ChIP assay. However, the one binding site effectively regulated the *oct4* promoter activity, as revealed in the zebrafish embryo luciferase assay (Figure 2T-2V). These observations suggest that Yy1 can directly regulate the promoters of RAGs positively or negatively.

### Yy1 overexpression positively regulates retinal progenitors

As the downregulation of Yy1 negatively affected the retinal progenitors during retina regeneration, we further explored the effect of its overexpression. For this we adopted an mRNA *in vivo* transfection strategy^44^. The overexpression of both *yy1a* and *yy1b* mRNAs profoundly increased the number of retinal progenitors (Figures 3A-3C). Yy1a overexpression caused a prominent increase in the number of retinal progenitors than Yy1b.

**Figure 3.**
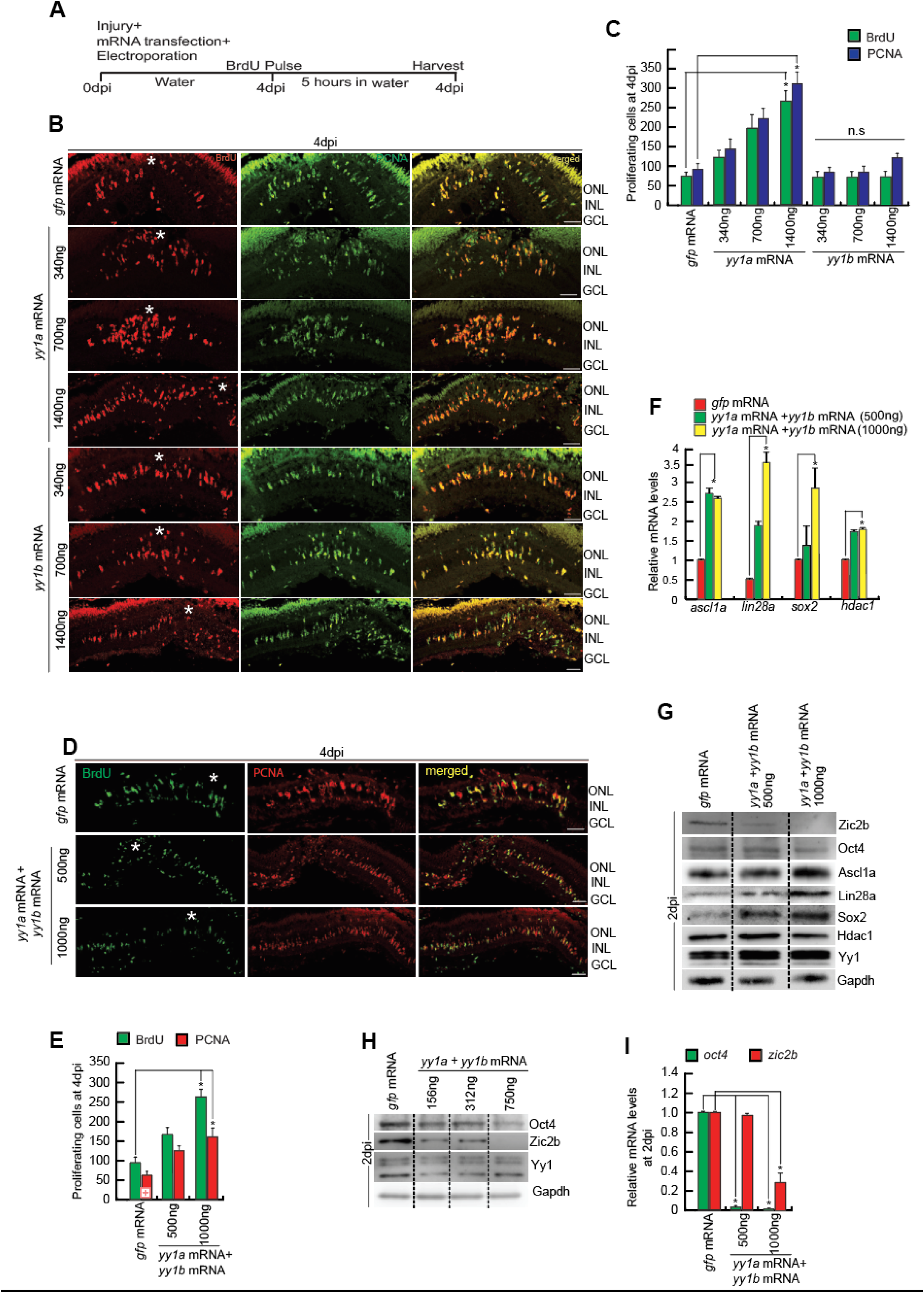
Overexpression of Yy1 increases the number of proliferating retinal progenitors post retinal injury. **(A)** An experimental timeline that describes injury, mRNA transfection and electroporation at 0dpi, BrdU pulse for 5hrs at 4dpi, followed by harvesting. **(B and C)** IF microscopy images of retinal cross-sections show a significant increase in the number of BrdU^+^ and PCNA^+^ retinal progenitors in *yy1a* overexpression, while no significant increase in the case of *yy1b* overexpression as compared to control *gfp* mRNA transfected retina at 4dpi (B), which is quantified (C). **(D and E)** IF microscopy images of retinal cross-sections show a significant increase in the number of BrdU^+^ and PCNA^+^ retinal progenitors in *yy1a* and *yy1b* combined overexpression, as compared to control *gfp* mRNA transfected retina at 4dpi (D), which is quantified (E). **(F)** qPCR analysis shows an upregulation in the levels of *ascl1a, lin28a, sox2* and *hdac1* in the combined overexpression of *yy1a* and *yy1b.* **(G)** Western blot analysis shows an upregulation in the levels of Ascl1a, Lin28a, Sox2 and Hdac1, while downregulation of Oct4 and Zic2b in the combined overexpression of *yy1a* and *yy1b.* **(H)** qPCR analysis shows a downregulation in the levels of *oct4* and *zic2b* in the combined overexpression of *yy1a* and *yy1b.* Scale bars represent 10μm in (B and D); asterisk marks the injury site and GCL, ganglion cell layer; INL, inner nuclear layer; ONL, outer nuclear layer in (B and D); dpi, days post injury. Error bars represent SD. *p < 0.005 in (E, F); *p < 0.0005 (C, H); n.s is non-significant. n = 6 biological replicates.

Combined overexpression of *yy1a* and *yy1b* mRNAs also caused an additive effect on the total number of retinal progenitors (Figures 3D and 3E). These observations support the idea that Yy1 is an essential positive regulator of retinal progenitors during regeneration. The question that remained was whether Yy1 influenced the retinal progenitors through RAGs. We analyzed a few selected RAGs at the mRNA and protein levels in Yy1a and Yy1b overexpressed conditions to address this possibility. We saw a dose-dependent increase in the levels of RAGs both at RNA and protein levels at 4dpi (Figures 3F and 3G). Of the RAGs we had tested in *yy1a* and *yy1b* knockdown, only Oct4 and Zic2b had shown an upregulation (Figure 2D), which showed an opposite trend because of Yy1a and Yy1b overexpression (Figures 3H and 3I). These findings indicate that the Yy1 protein has a differential effect on RAG regulations.

We explored if the increased number of retinal progenitors in the Yy1 overexpressed retina stayed back and contributed to the formation of retinal cell type. We performed a long-term experiment in which the injured retina transfected with *yy1a* and *yy1b* mRNA were given BrdU on 4, 5 and 6dpi. The fish recovered until 23 dpi, when the retina was harvested (Figure S3A). The retinal progenitor number, marked with BrdU, was substantially higher in 23dpi retina compared to *gfp* mRNA transfected control (Figures S3B and S3C), suggesting the survival of the increased number of retinal progenitors until 23dpi a time when retinal differentiation comes to a halt. We then explored the 23dpi retina to find if retinal progenitors formed in Yy1 overexpressed conditions could make various retinal cell types (Figure S3D). These observations support the idea that Yy1 overexpression causes an increase in the number of retinal progenitors that are viable and capable of differentiating into retinal cell types.

The morpholino-mediated knockdown of *yy1a* and *yy1b* resulted in a drastic decline in the retinal progenitor described in Figure 2. To prove that the effect of MOs was not because of their off-target effects, we performed a rescue experiment in which the *yy1a* and *yy1b* mRNAs with their MO-binding region mutated while retaining the same amino-acid codon were transfected in the *yy1a*/*yy1b* MO or control MO electroporated conditions. Retinal sections with *yy1a* or *yy1b* MO showed a drastic decline in the number of retinal progenitors, which restored to that of control-MO electroporated conditions upon co-transfection with *yy1a* and *yy1b* mRNAs respectively (Figure S3E and S3F). These results suggest that the reduced number of retinal progenitors in *yy1a*/*yy1b* knockdown could be rescued by *yy1a*/*yy1b* mRNAs. While the *yy1a* and *yy1b* mRNA transfected in the control-MO electroporated retina caused an increase in the retinal progenitors, interestingly, it had no appreciable effect when transfected into an uninjured retina (Figure S3G).

### Delta-Notch and Tgf-β signaling pathways affect Yy1 expression during retina regeneration

Delta-Notch signaling facilitates zebrafish retinal progenitor proliferation^45, 46^ unlike seen in mammals. We found upregulation in the levels of *yy1a* and *yy1b* in Delta-Notch signaling-blocked conditions through DAPT treatment (Figures 4A and 4B). Next, we checked the effect on the expression of *yy1a* and *yy1b* in the *her4.1*, the effector of Delta-Notch signaling, knockdown retina. The presence of the Hes/Her binding site in the promoter of *yy1a* and *yy1b* indicates their possible direct regulation through Delta-Notch signaling. In the *her4.1* overexpressed retina, we saw an expected downregulation of both *yy1a* and *yy1b* mRNAs (Figure 4C). The lack of Her4.1-mediated repression may be causing the upregulation of *yy1a* and *yy1b* in the DAPT-treated retina. We have previously shown the importance of Tgf-β signaling during zebrafish retina regeneration^13^. Unlike in mammalian systems, the Tgf-β signaling is pro-proliferative during zebrafish retina regeneration^3, 13^. During the early phase of zebrafish retina regeneration, the inhibition of Delta-Notch signaling upregulates Tgf-β signaling^13^. We explored whether the Tgf-β signaling affected the regulation of *yy1* transcription. We delivered Tgf-β1 protein into the injured zebrafish eye and monitored the expression levels of *yy1a* and *yy1b*. We saw a dose-dependent increase in the levels of Yy1 protein because of TGF-β1 protein delivery (Figure 4D). We also saw a dose-dependent decline in the *yy1a* and *yy1b* mRNA and Yy1 protein levels in Tgf-β signaling blocker SB431542-treated retina at 2dpi (Figures 4E and 4F). ChIP analysis of the 5GC elements on the yy1a promoter and yy1b first exon reveals that pSmad3 binds to these sites. (Figures 4G-4J). These observations support the idea that the *yy1* gene is positively regulated via Delta-Notch and Tgf-β signaling pathways.

**Figure 4.**
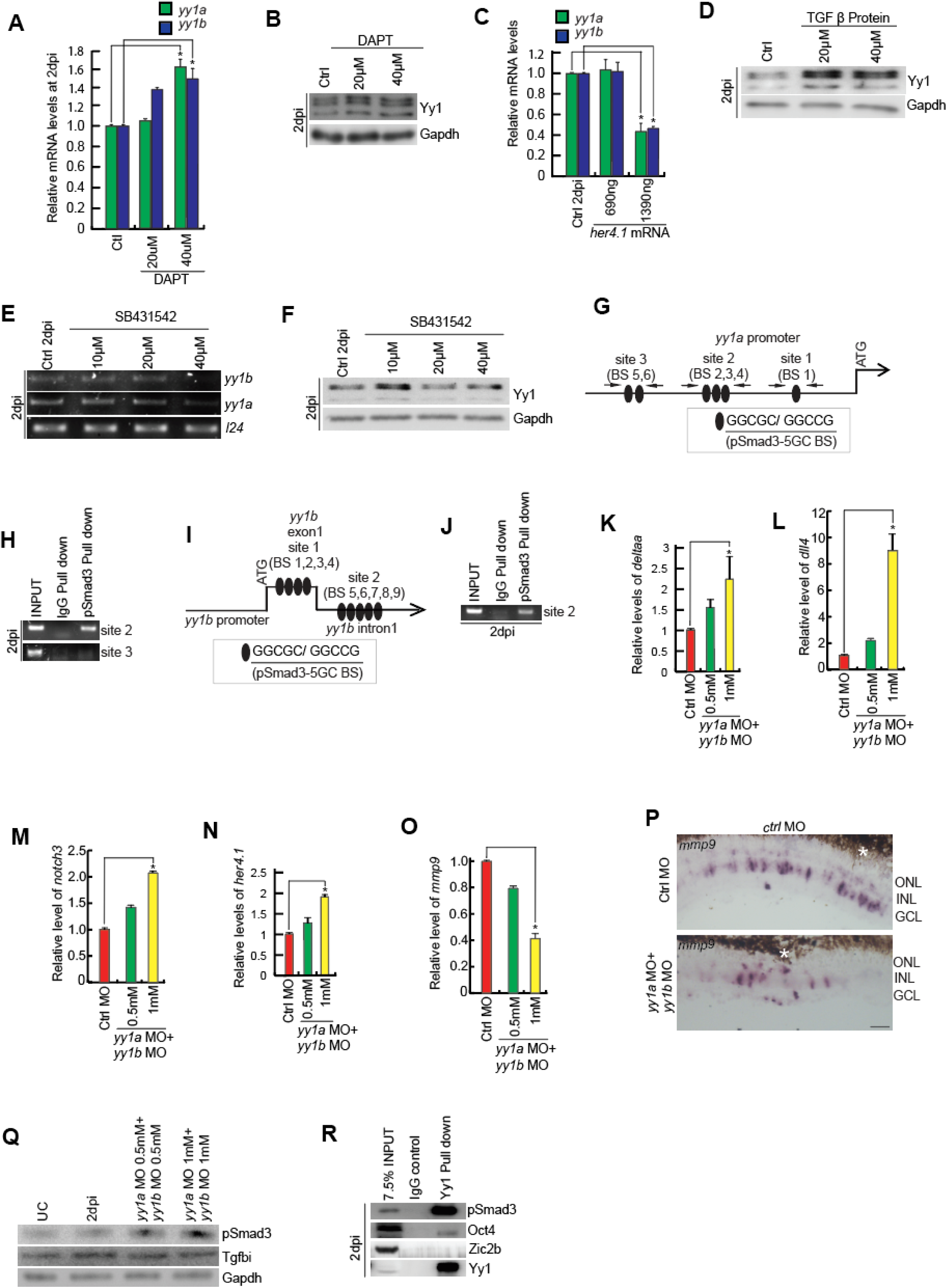
Yy1 is regulated by Delta-Notch and TGF-β signaling and in turn it also regulates Delta-Notch signaling components. **(A)** qPCR analysis shows an upregulation in the levels of *yy1a and yy1b* in a concentration-dependent manner in DAPT treatment. **(B)** Western blot analysis shows an upregulation in the levels of Yy1 in a concentration-dependent manner in DAPT treatment. **(C)** qPCR analysis shows decrease in the levels of *yy1a and yy1b* at the highest concentration of *her4.1* mRNA **(D)** Western blot analysis shows an upregulation in the levels of Yy1 in a concentration dependent manner in TGFβ protein treatment. **(E and F)** RT-PCR analysis of *yy1a* and *yy1b* (E) and western blot analysis of Yy1 (F) shows a decline in their levels in SB431542 drug treatment in a concentration-dependent manner. **(G and H)** The schematic showing the pSmad3 binding site (5GC-BS) on *yy1a* promoter (G) and the retinal ChIP assays confirm the physical binding of pSmad3 on *yy1a* promoter in 2dpi retina (H). **(I and J)** The schematic showing pSmad3 (5GC-BS) on *yy1b* 1^st^ exon and 1^st^ intron (I) and the retinal ChIP assays confirm the physical binding of pSmad3 on *yy1b* intron, in 2dpi retina (J). **(K, L and M)** qPCR analysis shows upregulation in the levels of Delta-Notch signaling components *delta* (K), *dll4* (L), and *notch3* (M) in *yy1a* and *yy1b* knockdown conditions. **(N)** qPCR analysis shows upregulation in the levels of *her4.1* in *yy1a* and *yy1b* knockdown conditions. **(O)** qPCR analysis shows downregulation in the levels of *mmp9* in *yy1a* and *yy1b* knockdown conditions. **(P)** mRNA ISH also shows a downregulation in the levels of *mmp9* in the *yy1a* and *yy1b* knockdown condition, as compared to the control. **(Q)** Western blot analysis shows the levels of pSmad3 and Tgfbi in the combined knockdown of *yy1a* and *yy1b,* at 2dpi. **(R)** Western blot analysis of Co-immunoprecipitated samples shows the strong interaction of Yy1 with pSmad3 but weak interaction with Oct4 and no interaction with Zic2b, at 2dpi. Scale bars represent 10μm in (P); asterisk marks the injury site and GCL, ganglion cell layer; INL, inner nuclear layer; ONL, outer nuclear layer in (P); dpi, days post injury. Error bars represent SD. *p < 0.001 in (A, C, K, L); *p < 0.005 in (M, N, O). n = 6 biological replicates. BS represents the binding site.

Interestingly, *yy1* negatively influenced the Delta-Notch signaling through the *delta, dll4, notch3,* and the effector *her4.1* expression levels (Figures 4K-4N). This regulatory loop suggests that the Delta-Notch pathway negatively influences *yy1a* and *yy1b* directly and via Tgf-β signaling. The Yy1, in turn, inhibits the components of the Notch signaling pathway, creating a yin-*yang* feedback loop on its expression. Furthermore, the *yy1* knockdown negatively affected the matrix metalloproteinase 9 (*mmp9*) levels (Figures 4O and 4P), which could also be because of elevated *her4.1* levels, a known repressor of *mmp9* during retina regeneration^12, 17^.

We then explored the levels of pSmad3 and Tgfbi, the effector and read out of Tgf-β signaling, respectively, in Yy1 downregulated retina using *yy1a* and *yy1b* MO (Figure 4Q). The mRNA levels of the components of Tgf-β signaling showed an increase in the *yy1a* and *yy1b* knockdown retina (Figure S4A), supporting the increased pSmad3 levels. Interestingly, we saw an increase in the pSmad3 levels and a steady level of Tgfbi, suggesting the possible ineffectiveness of the increased pSmad3 to upregulate Tgfbi. This made us speculate about the possible interaction of Yy1 and pSmad3 during genome targeting, as reported in other systems^47, 48^. We performed a co-immunoprecipitation of Yy1 in 2dpi retinal extracts and probed for pSmad3 alongside two other proteins Oct4 and Zic2b (Figure 4R). Oct4 and Zic2b are essential for normal retina regeneration, which gets upregulated post-retinal injury and in Yy1 knockdown conditions. We saw a good representation of pSmad3 in the Yy1 protein complex with negligible levels of Oct4 and Zic2b. We then performed another experiment where Tgf-β1 protein was injected intravitreally, which caused an increase in the retinal progenitor population^13^. Here, we speculated if the Yy1 was one of the causative factors for accelerated cellular proliferation in the Tgf-β1 protein injected scenario. We downregulated Yy1 levels in Tgf-β1 protein injected retina using *yy1a* and *yy1b* morpholinos and found that the number of retinal progenitors decreased significantly (Figures S4B and S4C), suggesting the possible dependence of Tgf-β signaling on Yy1 protein.

### Functionality of Yy1 with and without acetylation during retina regeneration

The Yy1 protein is known for its transcriptional activation and repression roles. It also collaborates with histone acetyltransferases such as p300 and PCAF, resulting in the acetylation of amino acid residues from 170-200 and the C-terminal region, essential for its transcriptional activation function^49^. The acetylated Yy1 could collaborate with HDAC1 to locate specific chromatin histone proteins to cause their deacetylation to ensure transcriptional repression. These HDACs also deacetylate Yy1, causing a negative feedback loop. It has been demonstrated that the regulation of Yy1 acts through its acetylation and deacetylation^26^ or phosphorylation of the DNA-binding domain^28^. An amino acid comparison between humans and zebrafish revealed five conserved lysine residues in Yy1a and four in Yy1b at the HAT/HDAC interacting domain (Figure S5). A previous study has demonstrated that mutation of these lysine residues to arginine abolishes the transcriptional repressive activity of Yy1^49^. In this scenario, we wished to mutate all six basic residues, including five lysine and one arginine residue of Yy1a and four from Yy1b to an acetylation mimetic mutation to glutamine, as successfully demonstrated for histone ^50, 51^ Tau ^52^and TRIM28^53^ proteins. The neutral mutation was lysine to alanine in respective positions. These *yy1a* and *yy1b* mRNAs with acetylation mimetic and neutral mutations were used to transfect the retina to explore the effect on retinal progenitor proliferation (Figure 5A). While the control GFP mRNA had no profound effect on the total number of retinal progenitors, the wild-type *yy1a* and *yy1b* RNA combination caused a pan-retinal progenitor induction (Figures 5B and 5C). Interestingly, both *yy1a* and *yy1b* mRNAs harboring neutral (K to A) or acetylation mimicking (K to Q) mutation had a significant decline in the retinal progenitors (Figures 5B and 5C). This observation suggests that the neutral and acetylation mimetic RNAs caused a dominant negative effect wherein neither Yy1 protein with neutral or acetylation mimetic could complement the full functionality of wild-type protein. This result suggests that the Yy1 protein would have a role in its acetylated and deacetylated form. It is interesting to note that several of the regeneration-associated proteins, such as Ascl1a, Sox2, Zic2b, Oct4, and Lin28, did not have a profound effect except for Oct4 and Zic2b, which showed an upregulation as seen with Yy1 knockdown (Figure 5D). To further confirm these results, we decided to block the HDAC activity, which will probably retain the Yy1 in its acetylated form in wild type and transfected mRNA, if eligible. In an experimental setup, we transfected the retina with *yy1a* and *yy1b* mRNA with or without Trichostatin A (TSA), a known blocker of HDACs (Figure 5E). We saw a significant decline in retinal progenitors when treated with TSA with or without *yy1a* and *yy1b* mRNA transfection when analyzed at 4dpi (Figures 5E and 5F). These results suggest that Yy1 could not perform its role in inducing adequate retinal progenitors in its acetylated form. It is important to note that the pan-retinal induction of retinal progenitors because of *yy1a* and *yy1b* overexpression also get nullified because of TSA treatment supporting the idea that Yy1 needs to undergo timely deacetylation during retina regeneration (Figures 5E and 5F). We then explored if the Yy1 protein interacted with Hdac1 during retina regeneration. We performed a co-immunoprecipitation assay on 2dpi retinal extract using a Yy1-specific antibody and probed for Hdac1 protein (Figure 5G). We saw a good intensity of the Hdac1 protein that got pulled along the Yy1 protein, suggesting the deacetylation of the Yy1 protein and the probable genome targeting of the Yy1-Hdac1 complex. When a few of the regeneration-associated proteins were analyzed, we found that despite an increase in the levels of Ascl1a, Lin28 and Sox2 compared to 2dpi control, the retinal progenitor number was on decline in Yy1 overexpressed with HDAC-inhibited retina at 4dpi suggesting the necessity of balanced acetylation and deacetylation of Yy1 protein during retina regeneration (Figure 5H). These observations support the idea that Yy1 collaborates with Hdac1 for its multitude of functions during retina regeneration.

**Figure 5.**
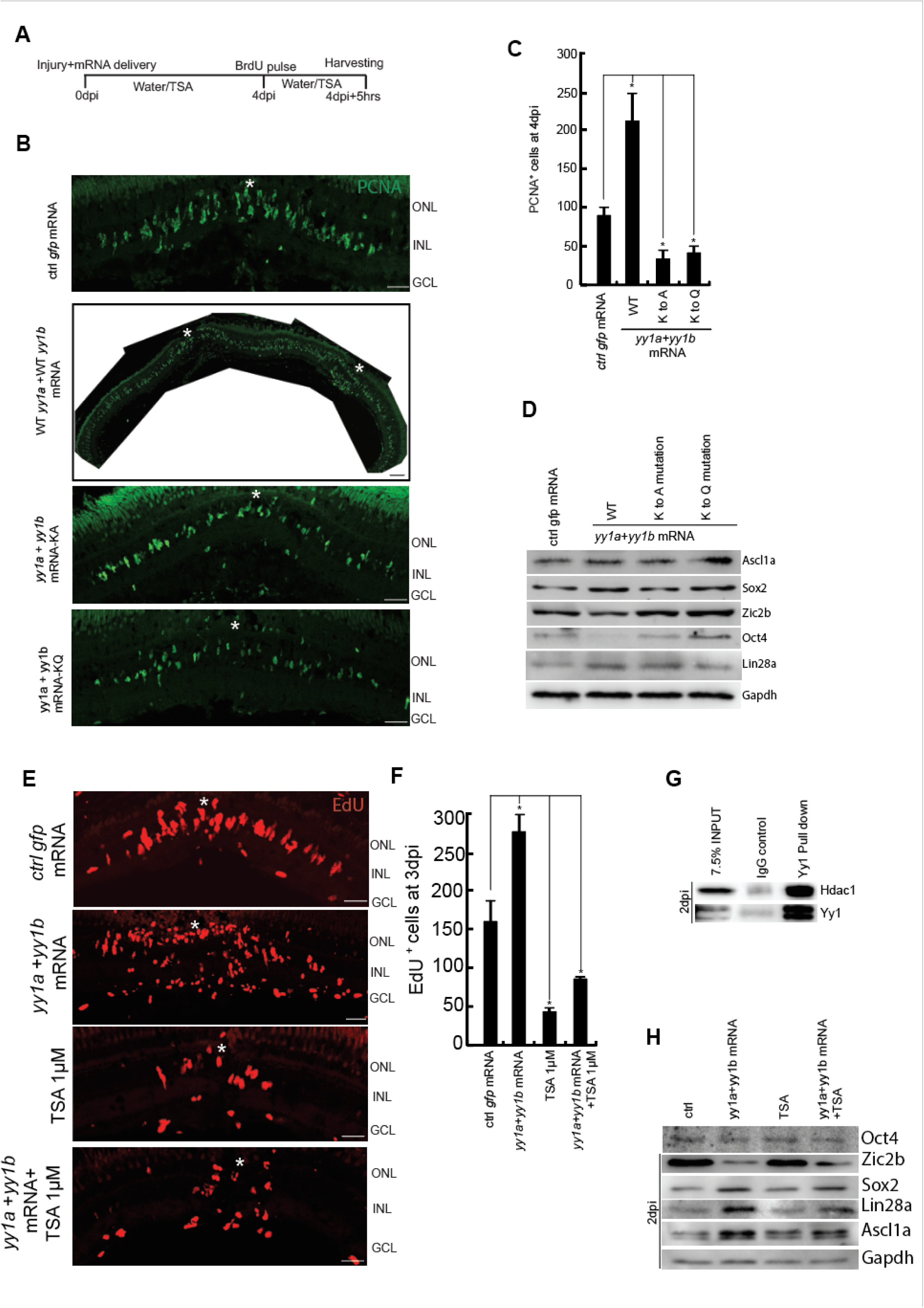
Regulation of activity of Yy1 by HATs and HDACs. **(A)** An experimental timeline that describes injury, wild-type and mutant mRNA delivery and electroporation at 0dpi, BrdU pulse for 5hrs at 4dpi, followed by harvesting. **(B and C)** IF microscopy images of retinal cross-sections at 4dpi, show an increase in the number of proliferating cells in the overexpression of wild type mRNA of *yy1a* and *yy1b*, which are marked by PCNA^+^ retinal progenitors, while decrease in the overexpression of both neutral mutation and acetylated-mimetic mutation (B), which is quantified also (C). **(D)** Western blot analysis shows the regulation of various RAGs in the experimental condition mentioned in B and C. **(E and F)** IF microscopy images of retinal cross-sections show an increase in the number of proliferating cells in the overexpression of wild type mRNA of *yy1a* and *yy1b*, which are marked by EdU^+^ retinal progenitors, while decrease in the TSA-mediated inhibition and in the combined treatment of TSA along with the overexpression of *yy1a* and *yy1b* (E), which is also quantified in (F). **(G)** Western blot analysis of co-immunoprecipitated samples shows the interaction of Yy1 with Hdac1, at 2dpi. **(H)** Western blot analysis shows the regulation of various RAGs in the experimental condition mentioned in E and F. Scale bars represent 10μm in (B and E); asterisk marks the injury site and GCL, ganglion cell layer; INL, inner nuclear layer; ONL, outer nuclear layer in (B and E); dpi, days post injury. Error bars represent SD. *p < 0.001 in (C) *p < 0.05 in (F). n = 6 biological replicates (C, D, F and H). KA refers to lysine to alanine mutation and KQ refers to lysine to glutamine mutation.

### Involvement of BAF complex in genome targeting of Yy1

Despite the earlier discovery of Yy1 to be associated with polycomb repressor complex 2 (PRC2) and its subsequent role as a transcriptional repressor ^54^, several recent studies revealed the importance of Yy1 as a transcriptional activator in collaboration with the BAF complex^55^. We explored the levels of a few important BAF complex gene family members, such as arid1aa, arid1ab, smarca2, and smarca4, at different stages after retinal injury (Figure 6A). Compared to the uninjured retina, we saw a declining trend for all these four genes from the 1-4dpi window, when retinal progenitors form and proliferate. This supports the view that similar to the declining levels of Yy1 (Figure 1A), its collaborator BAF complex also shows a similar expression pattern in the regenerating retina. We then explored whether we could block the BAF complex function using a pharmacological inhibitor, PFI-3. This drug targets the chromatin-bound BAF complex and prevents it from binding to DNA targets^56^. Exposure of the injured retina to increasing concentrations of PFI-3 caused a drastic decline in the number of retinal progenitors at 4 dpi (Figures 6B and 6C). As found with *yy1* knockdown (Figure 2D), or its pharmacological inhibition (Figure S2G), we saw an increase in the levels of *zic2b* and *oct4* levels with PFI-3 treatment (Figure 6D). Another transcriptional repressor and a facilitator of the retinal progenitors to exit the cell cycle ^57^, *insm1a*, also showed increased expression upon PFI-3 treatment (Figure 6D). On the contrary, *ascl1a*, which is expressed in the proliferating retinal progenitors ^19, 58^, and Hdac1, which is expressed in the neighboring cells of the retinal progenitors ^16^, did not show a significant change in the expression levels because of PFI-3 treatment (Figure 6E). These observations suggest that Yy1 and BAF complex show a parallel trend in their expression after a retinal injury, and inhibition of either could facilitate the upregulation of Zic2b and Oct4 with a reduction in retinal progenitors.

**Figure 6.**
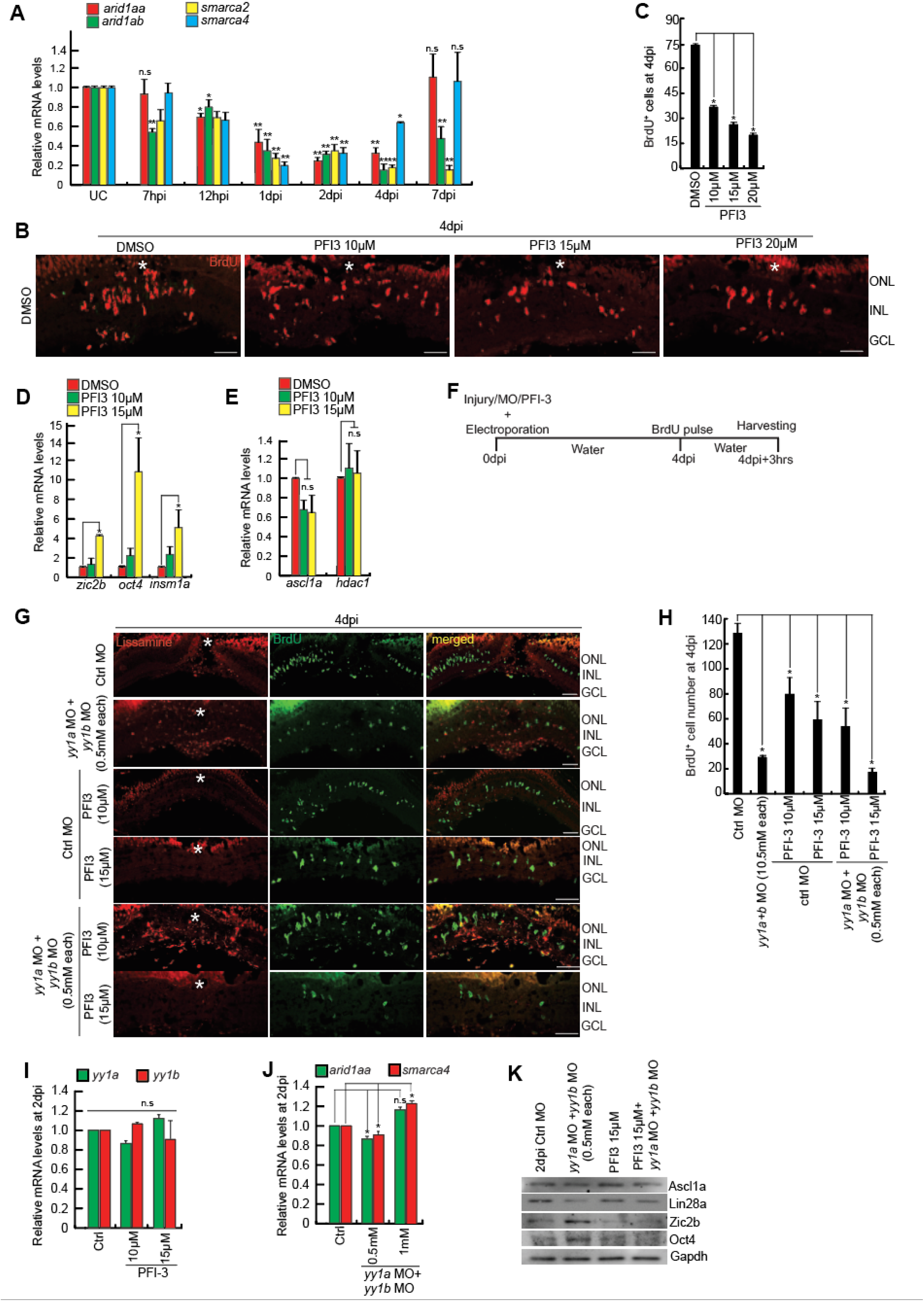
Yy1 and BAF complex have a synergistic effect on the proliferation of retinal progenitors post injury. **(A)** The qPCR analysis of BAF complex component genes, *arid1aa, arid1ab, smarca2* and *smarca4*, in the retina at various time points post-retinal injury. **(B and C)** IF microscopy images of retinal cross-sections at 4dpi, show a decrease in the number of proliferating cells in the PFI3 treatment, which are marked by BrdU^+^ retinal progenitors (B), which is quantified in (C). **(D and E)** qPCR analysis shows an upregulation in the levels of *zic2b*, *oct4* and *insm1a* (D), while no significant change in the levels of *ascl1a* and *hdac1* (E). **(F)** An experimental timeline that describes injury, mRNA delivery, dipping in the PFI3 drug and BrdU pulse for 5hrs at 4dpi, followed by harvesting. **(G and H)** IF microscopy images of retinal cross-sections at 4dpi, showed the combined effect of *yy1a* and *yy1b* knockdown and PFI3 treatment on the number of proliferating cells marked by BrdU (G), which is quantified in (H). **(I)** qPCR analysis shows that there is no significant change in the levels of both *yy1a* and *yy1b* in the retina of fish exposed to PFI3 drug. **(J)** qPCR analysis shows that there is no significant change in the levels of *arid1aa* and *smarca4* in the combined knockdown of *yy1a* and *yy1b*. **(K)** Western blot analysis shows the change in the levels of few of the RAGs in the experimental condition mentioned in G and H. Scale bars represent 10μm in (B and G); asterisk marks the injury site and GCL, ganglion cell layer; INL, inner nuclear layer; ONL, outer nuclear layer in (B and G); dpi, days post injury. Error bars represent SD. *p < 0.05 in (A), **p < 0.0001 in (A), *p < 0.001 in (C, D, E, H), *p < in (J); n.s is non-significant in (I). n = 6 biological replicates.

We then explored whether the Yy1 and BAF complex interact to maintain a healthy number of retinal progenitors in the regenerating retina. We performed *yy1a* and *yy1b* knockdown along with PFI-3 and evaluated the retina for progenitor cell number at 4dpi (Figures 6F-6H). We saw an additive effect in causing a decline in the retinal progenitors while using *yy1* MO and PFI-3, suggesting possible independent actions caused by Yy1 and BAF complex in regulating the number of progenitors during retina regeneration. Although the blocking of BAF complex did not affect the *yy1* levels (Figure 6I), there was a marginal increase in BAF complex component genes *arid1aa* and *smarca4* (Figure 6J) in 2dpi retina after *yy1a* and *yy1b* knockdown. When we explored the expression levels of a few regeneration-associated genes in 2dpi retina with BAF complex and Yy1-knockdown, either in isolation or in combination, there was an increase in the levels of Ascl1a, Lin28a and zic2b compared to control (Figure 6K). These observations support the idea that despite the increase in the expression of critical regeneration-associated genes, blocking either BAF or Yy1 negatively affects progenitor proliferation in an injured retina.

### Role of Yy1 in the absence of BAF complex

Our findings so far indicated the additive effect of Yy1 and BAF complex debilitation on the number of retinal progenitors, suggesting some independent roles during retinal regeneration. To test this, we blocked the BAF complex along with Yy1 overexpression (Figures 7A and 7B). Compared to the control *gfp* mRNA-transfected retina, the yy1a and yy1b mRNA transfection caused an approximate doubling of retinal progenitors (Figure 7C). The BAF complex inhibition caused an approximately 70% decline in the progenitor cell number (Figure 7C). Interestingly, the Yy1 overexpression rescued the reduction in progenitor cell number seen in the BAF complex-inhibited retina (Figure 7C). These observations support the idea that Yy1 and BAF complex contribute to the retinal progenitor number through independent pathways. We then explored the expression status of various regeneration-associated genes such as *ascl1a*, *lin28a*, *oct4*, *sox2*, *mmp9*, *her4.1,* and *hdac1* (Figure 7D). Compared to *gfp* mRNA transfected control, almost all genes except for *mmp9* and *hdac1* showed an upregulation in the retina because of Yy1 overexpression at 2dpi. Similarly, the blockade of the BAF complex decreased the expression of all except *oct4*, which showed a moderate increase. Interestingly, the overexpression of Yy1 along with BAF complex inhibition alleviated the decline in most cases, with a few reaching the levels to that of *gfp* mRNA transfected controls or even exceeding those levels. Notably, the *oct4* expression levels in BAF complex inhibited, and Yy1 overexpressed retina exceeded the levels seen either with Yy1 overexpression or with BAF complex inhibition alone, suggesting that BAF complex may have a profound negative effect on *oct4* expression compared to other genes. The mRNA levels of several of these genes are also reflected in their respective protein levels in 2dpi retina (Figure 7E). These observations suggested that despite inhibiting BAF complex function, the Yy1 overexpression drove the expression of several regeneration-associated genes supporting a BAF complex-independent pathway driven through Yy1 during retina regeneration.

**Figure 7.**
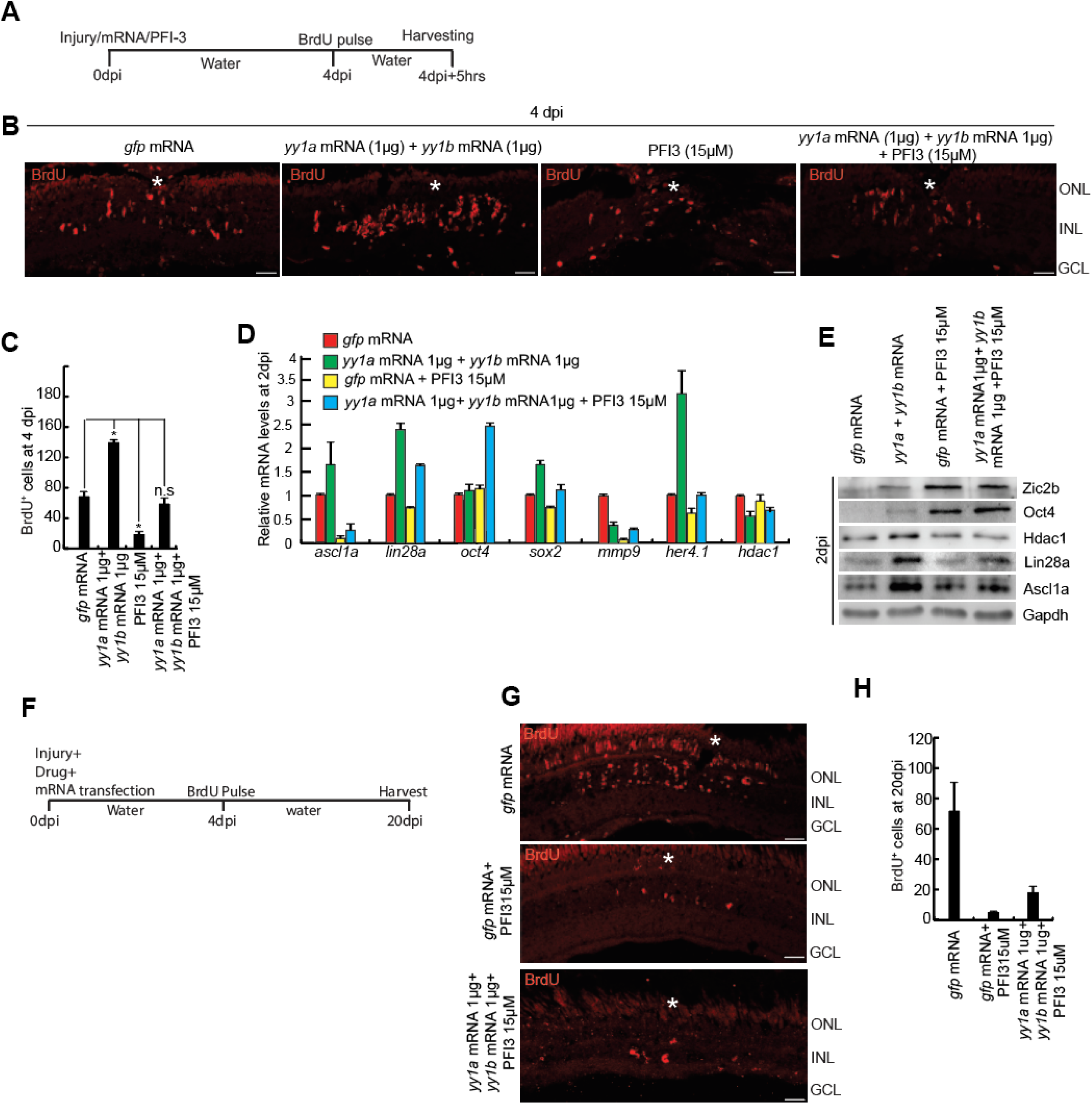
Yy1 requires BAF complex for the survival of proliferated cell post retinal injury. **(A)** An experimental timeline that describes injury, mRNA delivery, electroporation at 0dpi, dipping in the PFI3 drug and BrdU pulse for 5hrs at 4dpi, followed by harvesting. **(B and C)** IF microscopy images of retinal cross-sections at 4dpi, show the effect of combined treatment of PFI3 along with *yy1a* and *yy1b* overexpression on the number of proliferating cells, which are marked by BrdU^+^ (B), and quantified in (C). **(D)** qPCR analysis of few RAGs in the experimental conditions explained in B and C. **(E)** Western blot analysis shows regulation of Ascl1a, Lin28a, Hdac1 and Oct4 in the experimental conditions explained in B and C. **(F)** An experimental timeline that describes injury, mRNA delivery, electroporation at 0dpi, dipping in the PFI3 drug until 4dpi and BrdU pulse for 5hrs at 4dpi, followed by harvesting at 20dpi. **(G and H)** IF microscopy images of retinal cross-sections at 20dpi, shows the decrease in number of surviving BrdU positive cells, which were labelled at 4dpi in the combined treatment of PFI3 along with overexpression of *yy1a* and *yy1b* mRNA (G), which is quantified in (H). Scale bars represent 10μm in (B and G); asterisk marks the injury site and GCL, ganglion cell layer; INL, inner nuclear layer; ONL, outer nuclear layer in (B and G); dpi, days post injury. Error bars represent SD. *p < 0.001 in (C, D, H), n.s is non-significant. n = 6 biological replicates.

We were intrigued to know the fate of these retinal progenitors formed in the retina in the absence of BAF complex with or without Yy1 overexpression. For this, we adopted a long-term experimental regime in which retinal progenitors are labeled with BrdU at 4dpi, a time when retinal progenitor proliferation is at its peak, followed by retinal harvest at 20dpi, a time when retina regeneration is almost complete (Figure 7F). The retinal progenitors were labeled with BrdU at 4 dpi and followed up to 20 dpi in gfp mRNA-transfected controls, and BAF complex inhibited retina with or without Yy1 overexpression. The survival was more profound in the gfp mRNA-transfected controls than the BAF-inhibited ones (Figures 7G and 7H). These findings support the idea that despite the retinal progenitor proliferation at 4dpi, the absence of BAF complex renders them less viable towards the end of retinal regeneration, suggesting the importance of BAF complex and downstream gene regulations. We were intrigued to find the reasons for the absence of retinal progenitors at 20dpi. We speculated that a possible premature cell cycle exit would prevent perpetual cell division and represent a good number of cells at 20dpi, as found in the *gfp* mRNA transfected conditions. These findings support the idea that BAF complex function is essential, despite having an abundance of Yy1, for the survival of retinal progenitors.

### Effect of Yy1 downregulation on BMP signaling

We decided to explore the pan-retinal effect of Yy1 knockdown through RNA-Seq analysis. The *yy1*-knockdown retinae were analyzed for differential expression of mRNAs at 4dpi (Figure 8A). We saw several pathways governing cellular processes such as cellular proliferation, wound healing, and growth factor stimulation are upregulated because of Yy1 knockdown. Interestingly, the BMP signaling and regulation of T-cell activation pathways are represented multiple times in both upregulated and downregulated transcriptome groups. We were intrigued to explore the importance of Yy1 in regulating the BMP signaling. At first, we explored the importance of BMP signaling through its pharmacological inhibition. We used the drug K02288, which selectively blocks BMP-mediated Smad pathway activation without affecting TGF-β signaling^59^. Administering K02288 during retina regeneration significantly reduced the proliferation of retinal progenitors in a concentration-dependent manner (Figures 8B and 8C). We explored if the K02288 treatment was sufficient to downregulate BMP signaling through the expression levels of a known BMP-Smad1/5/8 target gene, *id1*^60, 61^. We saw approximately a 40% decrease in the id1 gene in the K02288-treated retina (Figure 8D). However, the knockdown of *yy1a* and *yy1b* caused up to 70% decrease in *id1* gene expression (Figure 8E), which is reflected in the 4dpi retina (Figure 8F), suggesting the potential roles of Yy1 in regulating BMP signaling. To confirm whether the *id1* gene was downregulated because of changes in *smad* 1/5/8 genes, we analyzed their expression levels in the *yy1* knockdown retina at 2dpi. We saw a *yy1a* and *yy1b* MO-dependent decline in the levels of *smad1 and smad8* (Figure 8G) and an increase in the *smad5* genes (Figure 8H), possibly explaining the decrease in *id1* levels. We then explored if the Yy1 knockdown affected the negative regulator of BMP signaling, *noggin3*^62^, whose inhibition is also important for normal tissue regeneration^63, 64^. Interestingly, the Yy1 knockdown enhanced the expression levels of *nog3* in a *yy1a*/*yy1b* MO dose-dependent manner at 2dpi (Figure 8I). These observations suggest the importance of normal levels of Yy1 protein in the effective orchestration of retina regeneration via BMP signaling.

**Figure 8.**
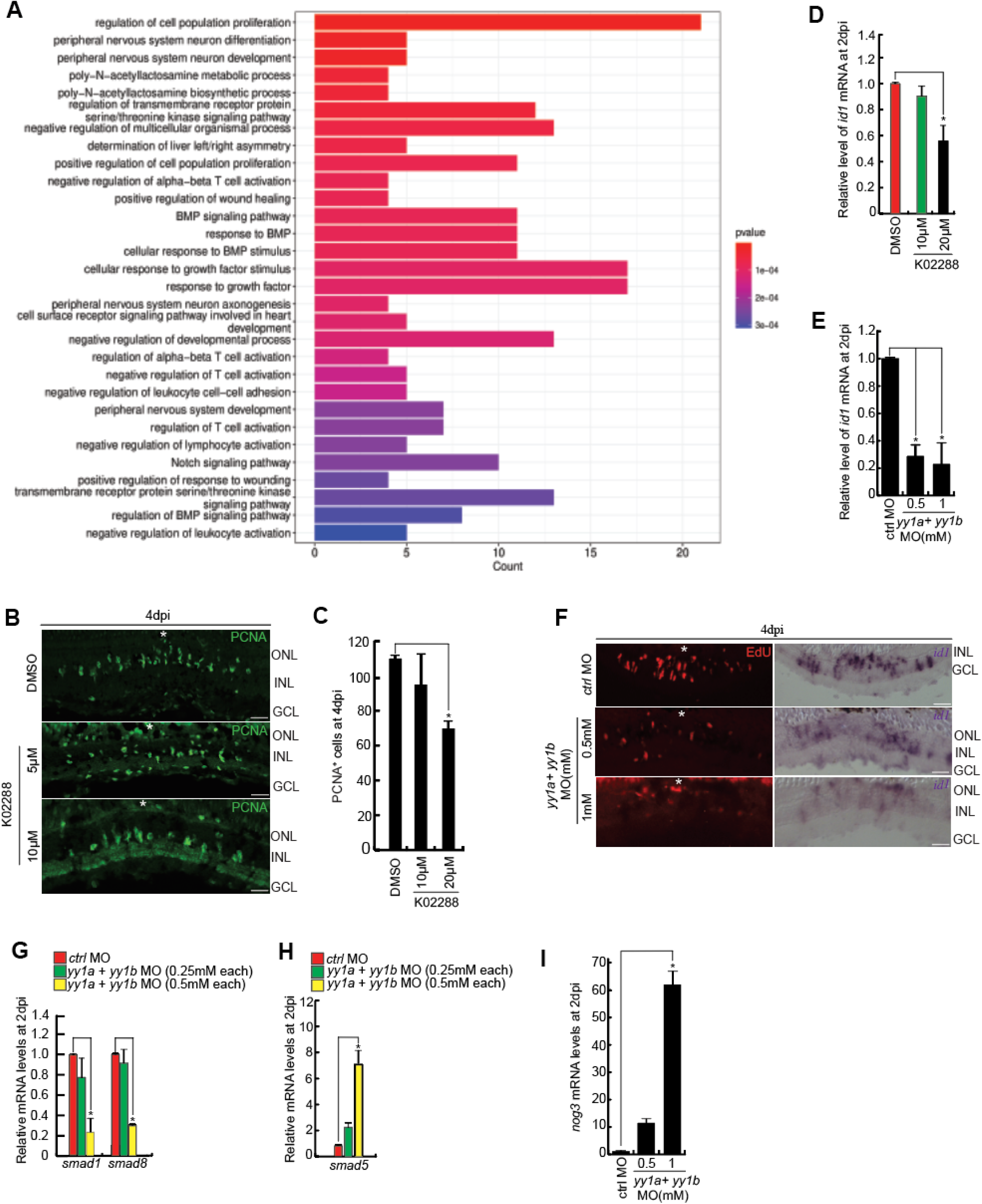
Yy1 regulates BMP signaling during retina regeneration and BMP signaling has a pro-proliferative role during retina regeneration. **(A)** GO function analysis of data from the whole retina RNA sequencing done in the knockdown of *yy1a* and *yy1b*, as compared to the control, shows downregulation of many genes related to cell proliferation, neuronal development and differentiation, wound healing, response to growth factors, Notch signalling and BMP signalling. **(B and C)** IF microscopy images of retinal cross-sections at 4dpi shows reduction in number of proliferating retinal progenitors, marked by PCNA, in the K02288 drug treatment (B), which is quantified in (C). **(D)** qPCR analysis shows decrease in the levels of *id1,* an effector gene of BMP signalling, in the K02288 drug treatment. **(E)** qPCR analysis shows decrease in the levels of *id1,* in the combined knockdown of *yy1a* and *yy1b.* (F) mRNA ISH shows decrease in the levels of *id1* at the injury site, in the combined knockdown of *yy1a* and *yy1b.* (G and H) qPCR analysis shows decrease in the levels of *smad1* and *smad8* (G) and an increase in the levels of *smad5* (H), in the combined knockdown of *yy1a* and *yy1b.* (I) qPCR analysis shows increase in the levels of *nog3*, a negative regulator of BMP signalling, in the combined knockdown of *yy1a* and *yy1b.* Scale bars represent 10μm in (B and F); the asterisk marks the injury site and GCL, ganglion cell layer; INL, inner nuclear layer; ONL, outer nuclear layer in (B and F); dpi, days post injury. Error bars represent SD, *p < 0.05 in (C, D, E, G, H), *p < 0.0001 in (I). n = 6 biological replicates.

## Discussion

The Yy1 is a very interesting protein with dual roles as a transcriptional activator or repressor. Yy1 undergoes acetylation and deacetylation in a context-dependent manner to cause unique gene expression events^26^. Yy1 is reported to play roles in axon^35^, spinal cord ^65^, and muscle regeneration^38, 39^. YY1 transcriptionally activates KLF4^66^, a well-known pluripotency-inducing factor. A recent study demonstrated the potential of Yy1 protein in collaboration with Ezh2 to drive the Hippo signaling effector Yap to cause gene repression events to promote cell proliferation^67^. YY1 is also implicated in long-distance DNA interactions^68^. Our study unravels the unique gene regulations and signaling pathways mediated via Yy1 during zebrafish retina regeneration. Despite surplus literature on the molecular dynamics and interactome of YY1, its role in tissue regeneration is limited^38, 39, 69, 70, 35, 41^. In the retina, the overexpression of Yy1 facilitates regenerative response in the injured retina but has no apparent roles in the uninjured retina, which suggest a need for wound response for Yy1 functionality.

The molecular association of Yy1 with acetyltransferase p300^71, 72^ and deacetylases such as Hdac1 is demonstrated as part of its gene targeting several systems. Yy1 is also functionally affected by its acetylation and deacetylation status^26^. During wound healing, the c-Fos-Yy1 complex collaborates with Hdac to cause repression of the PDGF receptor, which negatively influences wound healing^73, 74^. In this study, we demonstrated that the solely acetylated or deacetylated form of Yy1 is insufficient, but the context-dependent acetylation/ deacetylation cycle of Yy1 holds the key to efficient retina regeneration. It is important to note that YY1 is capable of interacting with histone acetyltransferase or histone deacetylases on IFN-beta gene regulatory sequence to cause activation and repression, respectively^75^, which support our observation that neither the acetylated nor the deacetylated Yy1 is sufficient during retina regeneration. During retina regeneration the Hdac1 levels dip soon after injury before restoring to normal levels, and staying secluded from the proliferating retinal progenitors^16^ which draws parallel with the Yy1 expression pattern characterized in this study. These attributes suggest the inevitable functional collaboration between Hdac1 and Yy1 during retina regeneration.

This study unraveled the importance of Yy1 during zebrafish retina regeneration through differential regulation of several regeneration-associated genes (RAG). Interestingly, important RAGs such as *ascl1a* and *lin28a* stay downregulated in *yy1* knockdown retina. The low Lin28a causes an abundance of *let-7* microRNA, the downregulation of which is essential during zebrafish retina regeneration^17, 19^. Yy1 downregulation positively regulated Delta-Notch signaling, reflected in the elevated *her4.1* levels, a transcriptional repressor and effector of Notch signaling. Her4.1 is a known repressor of Mmp9^17^, which could also contribute to its downregulation in the *yy1* knockdown retina.

We demonstrated that *yy1* gene expression is positively influenced by Tgf-β signaling, which is pro-proliferative during retina regeneration. This study also demonstrated the physical interaction of pSmad3 and Yy1 in regenerating retina. Similarly, the YY1-Smad1/2/3/4 interactions in the nucleus prevent the Smad proteins from finding selective gene targets as part of TGF-β or BMP signaling^48^. Another study demonstrated the YY1-Smad7 interaction, which is negatively regulated by TGF-β signaling, and Smad7 and YY1 act in concert to block TGF-β-induced gene transcription^76^. Smad7 also increases the YY1-Hdac1 interaction, facilitating transcription repression^76^. Our study unraveled the interaction of Yy1 and pSmad3, which seems essential for effectively relaying Tgf-β signaling. This Yy1-pSmad3 interaction could account for accelerated progenitor proliferation upon Yy1 overexpressed regenerating retina. Single-cell RNA sequencing studies have revealed the importance of BMP-signaling and its effector *id1* expressed in reactive Müller glial cells of lesioned zebrafish retina, indicating its necessity during regeneration^77^. We saw a positive regulation of BMP-signaling components *smad1/5* and effector *id1* by Yy1, leading to retinal progenitor proliferation. A similar positive correlation between BMP signaling and retinal progenitors is seen during chick retina regeneration^78^. Furthermore, the Yy1, induced by Tgf-β/BMP signaling, could also act as a mediator to restrict the Tgf-β/BMP signaling-induced selective gene expressions either through sequestering Smads or by recruiting Hdac1 to various genome targets.

We saw two regeneration-associated genes, oct4 and zic2b, get upregulated because of Yy1 knockdown, suggesting a repressive role of Yy1 on these gene expressions. In zebrafish, the Beta-arrestin1 collaborates with Yy1 in preventing the latter from its ability to recruit polycomb group (PcG) to *cdx4* promoter leading to Cdx upregulation and its downstream hox genes^79^. Similarly, the protein Gon4l forms a complex with YY1, transcriptional repressor Sin3a, and Hdac1 to target DNA, causing gene repressions^80^. Competitive binding by Seryl-tRNA synthetase/YY1 complex and NFKB1 negatively regulated *vegfa* transcription^81^. During retina regeneration, the Yy1 negatively regulated Zic2b and Oct4 through its physical binding onto respective promoters, and the majority of the retinal progenitor-neighboring cells express Yy1 probably to repress these RAGs during cell cycle exit.

In this study, we also demonstrated the importance of the BAF complex in the transcriptional activation function of Yy1. Normal activity of the BAF complex is also essential for retinal progenitor proliferation. Overexpression of Yy1 can rescue the phenotype observed in BAF complex inactivation, but the survival of these progenitors seems bleak in the later stages of regeneration. The BAF complex needed for cell survival is demonstrated during murine neural crest development^82^. The mammalian SWI/SNF, also known as the BAF complex, is shown to be essential to prevent cellular apoptosis^83^, which causes the lack of progenitor survival when the BAF complex is inhibited during retina regeneration.

YY1 positively regulated the expression of SOCS3 to cause neuroinflammation^84^. YY1 along with GATA-4 form a transcriptional activator complex of B-type natriuretic peptide (BNP) gene expression^85^. The ability of YY1 to activate the vimentin gene is prevented if the CpG island near the transcription start site is methylated^86^. These attributes make the Yy1 a very complex transcriptional activator. In this study, we saw the physical binding of Yy1 onto regeneration-associated genes ascl1a, lin28a, and *yy1* itself. Yy1 seems to be a transcriptional activator on *ascl1a*, *lin28,* and *yy1* genes. A comprehensive model made from the results of this study is depicted in Figure S6.

## Methods

### Animal maintenance and breeding

Zebrafish were maintained at 28°C in an automated recirculatory system on a 14:10 hours light/dark cycle. Embryos were obtained from the natural breeding and were also maintained at 28°C in an incubator. Transgenic lines, *1016 tuba1a*: GFP was also maintained in the same conditions as the wild type.

### Retinal Injury, pharmacological inhibitors, morpholino injections, mRNA transfection and rescue experiments

Zebrafish, 6-12 months old, were anesthetized using tricaine methane sulphonate. Retina was stabbed for a minimum of 4 times for all the experiments, from the back of the eye using 30 G needle. Needle was inserted to the length of the bevel and fish were then left in water for the recovery ^87^.

The pharmacological inhibitors used in the study were either delivered through intra-vitreal injection using Hamilton syringe of 10μl capacity and 30gauge needle or by dipping the fish in the drug post retinal injury. All the drugs used in the study are: DETANONOate (Sigma Aldrich, cat number D5431), PFI3 (Sigma Aldrich, cat number SML 0939), K02288 (Tocris, cat number 4986), DAPT (Sigma Aldrich, cat number, D5942), SB431542 (Sigma Aldrich, cat number S4317), and Trichostatin A (TSA) (ApexBio, cat number 8183). Working concentration of all the drugs was made in autoclaved deionized water.

Lissamine-tagged morpholino (MOs) (Gene Tools) of approximately 0.5μl (0.5-1mM) was injected into the vitreous of the eye from one of the injury spots using Hamilton syringe with 30G needle. Their delivery into the cell was ensured by electroporation with five pulses at 70 V for 50 ms, as described previously ^88^. The control MO is same as used in previous studies and their sequence has been mentioned elsewhere ^45^. Sequences of *yy1a* and *yy1b* MO used in this study are as follows:

*yy1a* MO *–* CCATTCTTGGCTTTCTTGCTTTCCG

*yy1b* MO *-* TCTCCCGGACGCCATCGTTAA

For overexpression experiments, coding sequence of various genes were cloned in the pCS2^+^ Vector. They were then linearized at C-terminal and capped mRNA were synthesized *in-vitro* using mMESSAGE MACHINE SP6 kit (Ambion). Transfection mixture was prepared as described previously ^14, 16^ and injected into the vitreous and electroporated to increase the efficiency of mRNA delivery.

*In-vivo* rescue experiments were performed to check the specificity of the *yy1a* and *yy1b* morpholino. MO binding sites were mutated to create silent mutations and *in-vitro* synthesized mRNA was electroporated along with the MO and phenotypic effect on proliferation was seen through immunostaining. For confirming the efficient transfection, *gfp* mRNA was transfected in the control injured retina.

### Primers, plasmid construction and reporter line

All the primers used in the study are listed in the Table S1. The CDS of *yy1a* and *yy1b* were amplified from the 24hpf embryos, digested with the restriction enzyme and cloned in the pCS2^+^ vector.

The *ascl1a:gfp-luciferase, lin28a:gfp-luciferase, insm1a:gfp-luciferase*, *oct4:gfp-luciferase* and *zic2b:gfp-luciferase* constructs were described previously (^19^; ^57^ ^14^ ^17^.

The *1016 tuba1a*:GFP transgenic fish used in this study have been characterized previously (Fausett and Goldman, 2006).

### Total RNA isolation, RT-PCR and qPCR analysis

Total RNA was isolated from the dark adapted zebrafish retinae of the control, MO injected, drug-treated and mRNA transfected experimental sets using TRIzol (Sigma). The double stranded cDNA was reverse transcribed from 5μg of the total RNA using combination of random hexamers and oligo dT primers from cDNA synthesis kit (Thermo Fisher Scientific). Quantitative real time PCR was done with atleast six biological replicates and reaction was put in triplicate for each sample using BIORAD SYBR mix in Applied biosciences real time PCR detection system. The *let-7a miRNA* levels, in control versus treated condition, were assessed with TaqMan *hsa-let7-a* probe (Applied Biosystems) as per the manufacturer’s instructions. The relative expression of genes in control versus treated retinae was deciphered using the ΔΔCt method and normalized to β*-actin* or *l24* mRNA levels.

### CO-Immunoprecipitation (CO-IP) and western blot assay

For CO-IP, approximately 20 retinae were dissected after dark adaptation and frozen in C-100 lysis buffer ^89^. After thawing in ice chilled water, lysate was prepared by pipetting and thawing. Lysate was then centrifuged at high speed of 9425g for 20 minutes and supernatant was separated. Input sample was aliquoted from the supernatant and the remaining lysate was divided into two separate tube and diluted with the same buffer for efficient rotation. 2μl of Anti-Yy1 antibody was added to one tube and Rabbit IgG was added to the other and kept overnight for rotation at 4°C. Next day, Protein A beads were added and again kept for rotation for 2 hours. Then, beads were washed 5 times with the C-100 buffer and protein was eluted in 2X Laemmli buffer. Eluted sample along with the input were subjected to western blot assay as described below.

Western blotting was performed using single retina per experimental sample lysed in Laemmli buffer, size-fractioned in 12% acrylamide gel at denaturing conditions, and transferred onto Immuno-Blot polyvinylidene fluoride (PVDF) membrane (cat. no. 162-0177; Bio-Rad), followed by probing with specific primary antibodies and HRP-conjugated secondary antibodies for chemiluminescence assay using Clarity Western ECL (cat. no. 170-5061; Bio-Rad). Primary antibodies used in the study are Rb Anti-YY1, rabbit polyclonal antibody against human ASCL1/MASH1 (Abcam, cat. no. ab74065), Rabbit polyclonal antibody against LIN28a (Cell Signalling Technologies, cat. no. 3978), mouse monoclonal against Oct3/4 (sc5279; Santa Cruz Biotechnology), Mouse polyclonal antibody against Zic2b (raised in house in mice against full length zebrafish Zic2b protein as antigen), rabbit polyclonal antibody against Sox2 (cat. no. ab59776; Abcam), Rabbit polyclonal antibody against pSmad3 (Abcam, ab52903), Rabbit monoclonal antibody against Tgfbi (Abcam, ab170874). Secondary antibody used were described previously ^17^.

### Chromatin immunoprecipitation (ChIP) assay

ChIP assay was performed with 20 retinae, harvested after dark adaptation at 2dpi. Chromatin was isolated as described previously ^90^. After sonication, a part of supernatant was kept as input and remaining was divided into two halves. One half was pulled down with anti-Yy1 antibody (described above), while the other half was pulled down with Rabbit IgG (Sigma Aldrich, I5006) as negative control. Primers used for ChIP assays are described in Supplementary Table S1.

### Embryos microinjections and luciferase assay

For luciferase assays, single-cell zebrafish embryos were injected with a total volume of ∼1nl solution containing 0.02pg of *Renilla* luciferase mRNA (normalization), 5pg of *promoter:* GFP–luciferase vector and 0–6pg of *yy1a and yy1b* mRNA or 0.1–0.5 mM *yy1a and yy1b* MO. To assure consistency of results, a master mix was made for daily injections and ∼300 embryos were injected at the single cell stage. After 24h, the embryos were divided into three groups (∼70 embryos/group) and lysed for dual-luciferase reporter assays (E1910; Promega).

### Brdu and Edu pulsing, Immunostaining, mRNA *in-situ* hybridization and TUNEL assay

BrdU labeling was performed by a single intraperitoneal injection of 20μl of BrdU (10 mM) 5 h before euthanasia and retina dissection, unless mentioned specifically. EdU labeling was done by intravitreal injection of 10 mM EdU solution as described earlier ^15^ ^16^. Fish were given a higher dose of tricaine methane-sulphonate and eyes were dissected, lens removed, fixed in 4% paraformaldehyde, serially washed with sucrose and sectioned as described previously ^87^.

Immunofluorescence protocols and antibodies were previously described (Mitra et al., 2019;^91^. Some of the antibodies specifically used for this paper are Rb Anti-YY1 and Rb Anti PH3. EdU staining was performed with Click-iT EdU Alexa Fluor 647 Imaging kit (Thermo Fischer Scientific, C10085), following manufacturer’s instructions.

The mRNA in situ hybridization (ISH) was performed on 12μM retinal sections with fluorescein or digoxigenin-labelled complementary RNA probes (FL/DIG RNA labelling kit, Roche Diagnostics), as described previously ^92^).

TUNEL assay was performed on retinal sections using In Situ Cell death Detection Fluorescein kit (Roche, CAT number 11684795910) as per manufacturer-recommended protocol.

### Fluorescence and confocal microscopy, cell counting, and statistical analysis

All the slides obtained after immunostaining, mRNA *in-situ* hybridization and TUNEL, were mounted and examined with a Nikon N*i*-E fluorescence microscope equipped with fluorescence optics and Nikon A1 confocal imaging system. The PCNA^+^ and BrdU^+^ cells were counted manually by observing fluorescence and ISH signals in the bright field under the microscope. Observed data were analysed for statistical significance by comparison done using two-tailed unpaired *t* test in case of comparison between two groups. In case of comparison between more than two groups, analysis of variance (ANOVA) was performed followed by Dunnett test for post hoc comparison using graph pad prism. Error bars represent standard deviation (s.d.) in all histograms.

### Fluorescence based cell sorting

GFP^+^ Müller glia derived progenitor cells were sorted separately from GFP^-^ fraction of the retinae using FACS (Fluorescence-assisted cell sorting) at 4dpi, as described previously ^58^ ^57^. Briefly, uninjured and injured retinae were isolated from *1016tuba1a:*GFP transgenic fish. GFP^+^ MGPCs from *1016tuba1a:*GFP retinae at 4 dpi were isolated by treating retinae with hyaluronidase and trypsin and then sorted on a BD FACS Aria Fusion high-speed cell sorter. Approximately 20 injured retinae with 10 pokes per retina from *1016tuba1a:*GFP fish yielded 60,000 GFP^+^ and 150,000 GFP^−^ cells. Total RNA was isolated from both the fractions and expression levels of genes was quantified by real time PCR.

### Whole retina RNA seq and data analysis

Using bulk RNA-seq gene expression profile in *yy1a* and *yy1b* MO injected retinae was compared to control MO injected retinae at 4dpi. Briefly raw reads generated were quality trimmed and adapter sequences were clipped using Trimmomatic ^93^. Fast QC (Babraham Institute) ^94^ was used for visualizing read quality. For subsequent analysis, data was uploaded to the Galaxy web platform, and public server at usegalaxy.org was used to analyse the data^95^. The filtered reads were mapped to GRCz11/danRer11 genome assembly using STAR (RNA STAR) ^96^. The number of read mapping to each feature was quantified using HTseq-count ^97^. FPKM (fragments per kilobase of exon per million) values were calculated using raw counts of genes. Fold change in expression was calculated as compared to the control sample. Genes with >1.5-fold change were considered for subsequent analysis. Gene Ontology (GO) enrichment analysis was performed using R package “cluster profiler” ^98^.

## Acknowledgments

M.C. acknowledges her support from the IISER Mohali for Junior and Senior Research Fellowship. P.S. acknowledges postdoctoral fellowship support from DBT, RA (DBT-RA/2023/July/N/4351) and NPDF (PDF/2019/001148), DST India. S.P. acknowledges support from CSIR for Senior Research Fellowship. P.Shukla acknowledges CSIR Senior Research Fellowship. O.M.D acknowledges support from IISER Mohali. R.R also acknowledges research funding from STARS, DoE (MHRD) Government of India (STARS/APR2019/BS /180/FS), DBT India (BT/PR36570/BRB/10/1976/2021) and support from IISER Mohali. We also acknowledge the contribution of Mr. Prateek Arora for operating the BD FACS Aria Fusion high-speed cell sorter for sorting GFP^+^ MGPCs from *1016tuba1a*:GFP transgenic fish retina.

## Supplementary Figure legends

**Figure S1.**
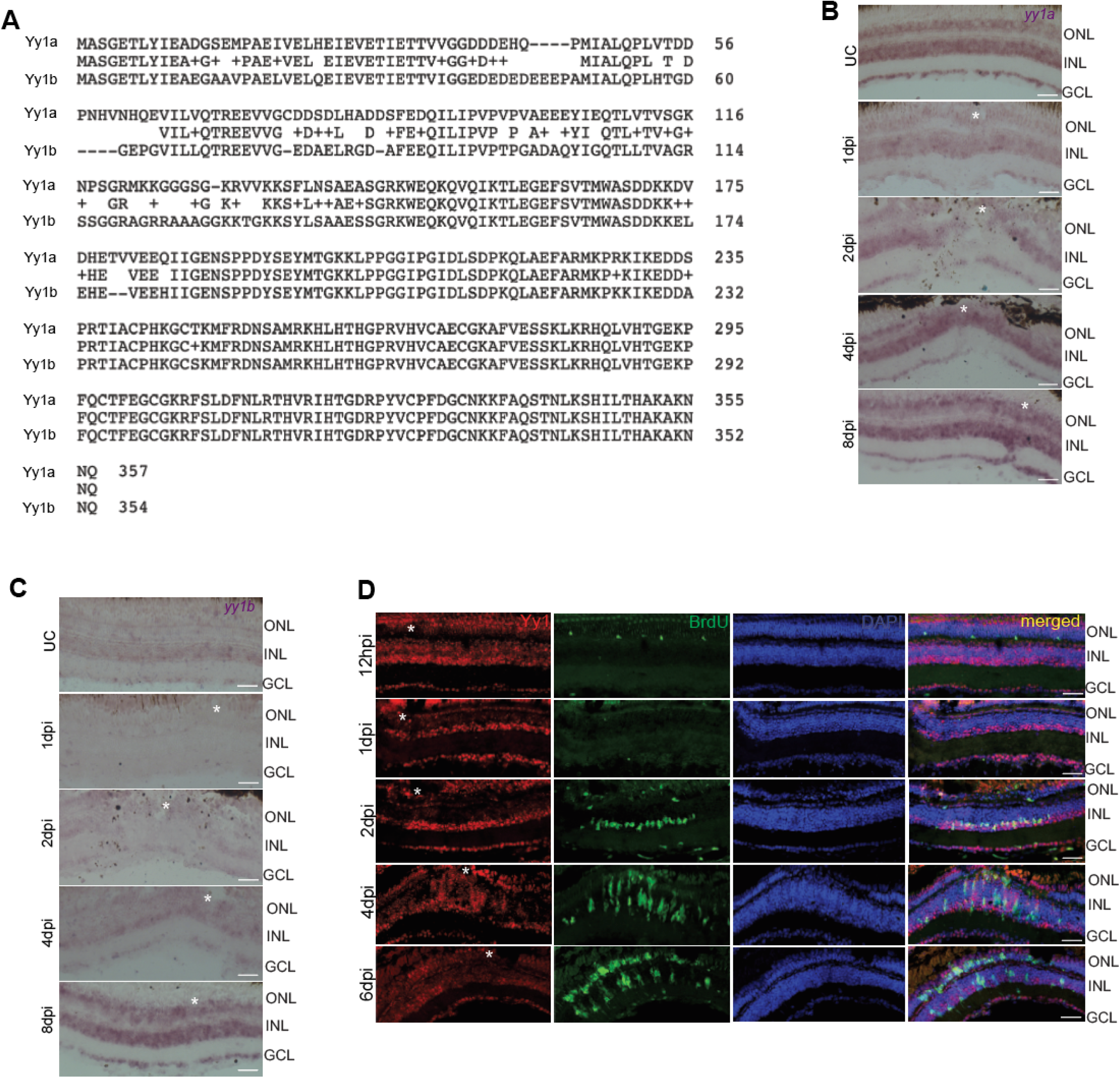
Sequence similarity between Yy1a andYy1b, Spatial time course analysis of *yy1a* and *yy1b* through mRNA *in-situ* hybridization. (**A**) NCBI BLAST analysis shows the amino acid sequence alignment between zebrafish Yy1a and Yy1b proteins. **(B and C)** 20X images of mRNA *in-situ* hybridization shows the spatial expression of *yy1a* (**B)** and *yy1b* (C) at different time-points post-retinal injury. **(D)** IF microscopy images of retinal cross-sections shows expression of Yy1at different time points post-retinal injury. Scale bars represent 10μm in (B, C, D); the asterisk marks the injury site and GCL, ganglion cell layer; INL, inner nuclear layer; ONL, outer nuclear layer in (B, C, D); dpi, days post injury; hpi, hours post injury.

**Figure S2.**
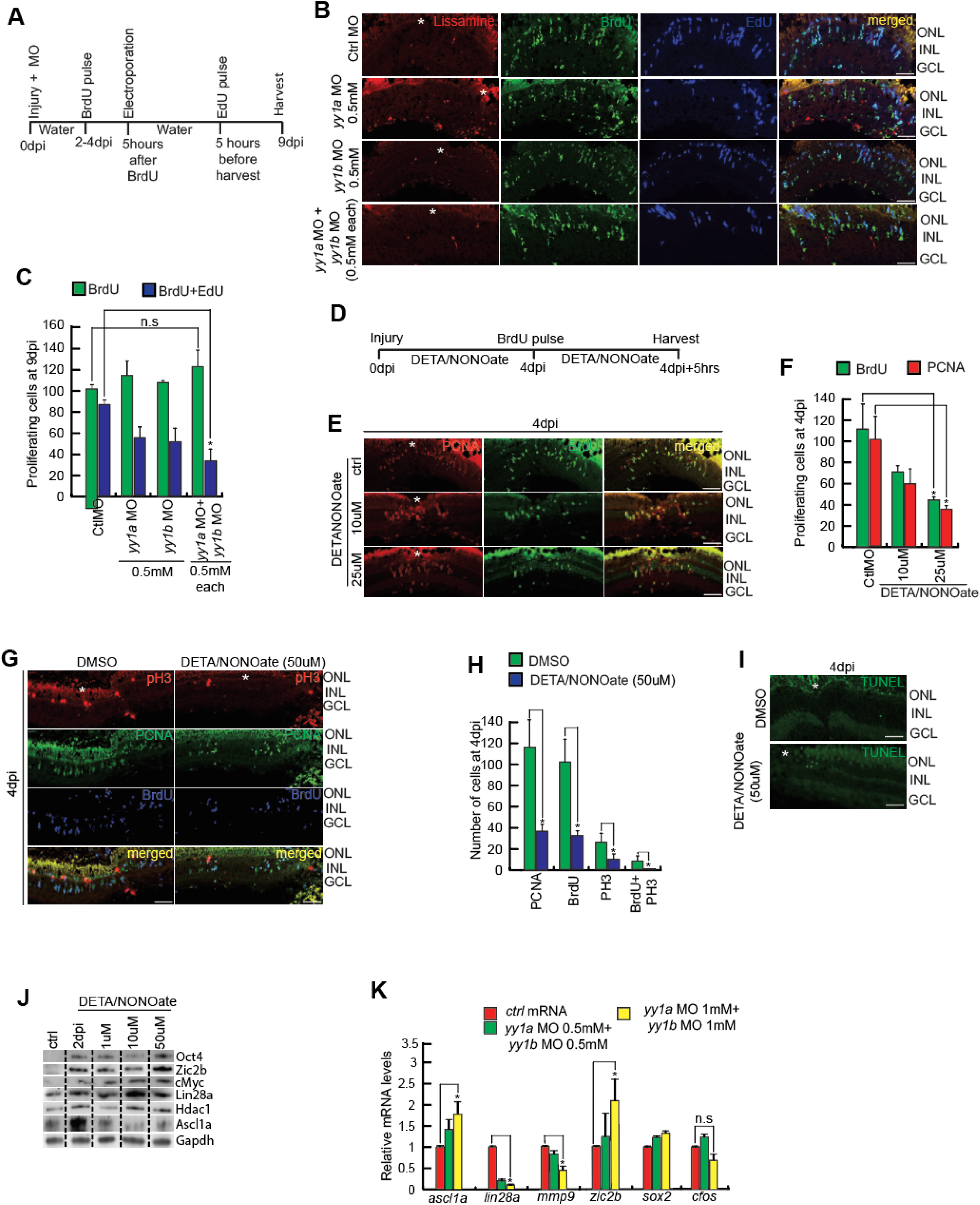
Late knockdown of Yy1 causes the early cell cycle exit of proliferating cells, and Yy1 inhibitor, DETA/NONOate, decreases the number of proliferating cells post retinal injury. **(A)** An experimental timeline that describes injury, MO delivery in the vitreous at the time of injury, BrdU pulse from 2-4dpi, electroporation at 4dpi and harvest at 9dpi post 5h of EdU pulse. **(B and C)** IF microscopy images of retinal cross-sections at 9dpi shows reduction in the number of EdU^+^ cells in the combined late knockdown of *yy1a* and *yy1b*, as compared to control (B), which is quantified also (C). **(D)** An experimental timeline that describes injury, dipping in the drug DETANONOate, BrdU pulse at 4dpi for 5 hours prior to harvest. **(E and F)** IF microscopy images of retinal cross-sections at 4dpi shows reduction in number of proliferating cells, as compared to control, which are marked by PCNA and BrdU (E), and quantified in (F). **(G and H)** IF microscopy images of retinal cross-sections at 4dpi shows decrease in the number of mitotic cells upon treatment with DETANONOate drug, which are marked by PH3 (G) and quantified in (H). **(I)** IF microscopy images of retinal sections at 4dpi shows no significant number of TUNEL^+^ cells in DETANONOate treatment. **(J)** Western blot analysis shows effect of DETANONOate on few RAGs in a concentration dependent manner at 2dpi. **(K)** qPCR analysis shows regulation of RAGs in the combined knockdown of *yy1a* and *yy1b*, at 2dpi. Scale bars represent 10μm in (B, E, G, I); the asterisk marks the injury site and GCL, ganglion cell layer; INL, inner nuclear layer; ONL, outer nuclear layer in (B, E, G, I); dpi, days post injury. * p<0.05 in (C), p < 0.0001 in (F and H)), p<0.05 in (K); n.s. non-significant.

**Figure S3.**
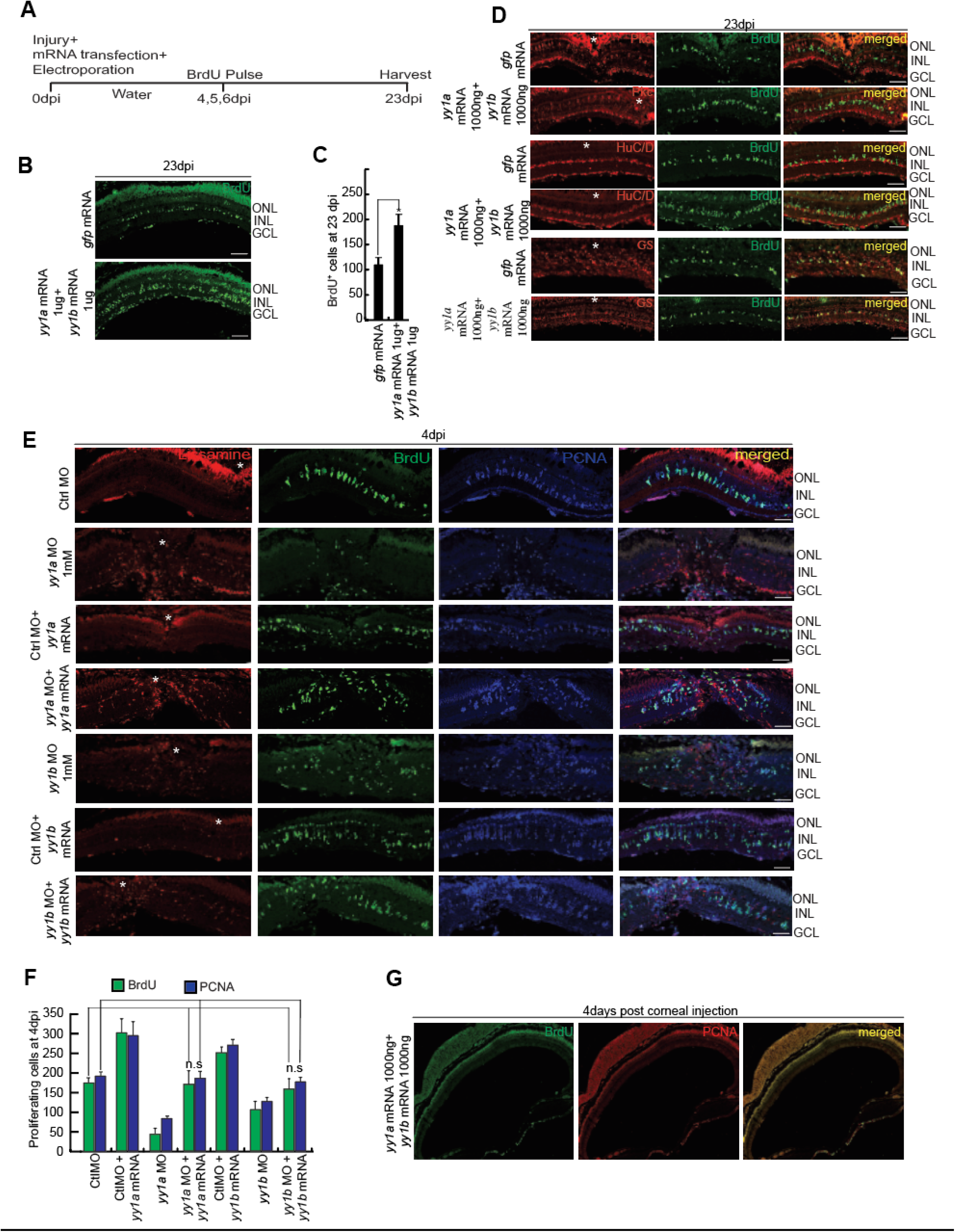
Lineage tracing of proliferating cells shows the survival of cells and their re-differentiation into various retinal cell types, and *yy1a* and *yy1b* MO-BS mutated mRNA could rescue the effect of *yy1a* and *yy1b* MO, respectively. (**A**) An experimental timeline that describes injury, mRNA delivery at the time of injury, BrdU pulse from 4-6dpi and harvest at 23dpi. **(B and C)** IF microscopy images of retinal section at 23dpi, shows an increased in the number and survival of BrdU^+^ retinal progenitors in the combined overexpression of *yy1a* and *yy1b* (B) and which is quantified also (C). **(D)** Long regime experiment mentioned in (A), shows that the increased number of BrdU^+^ retinal progenitors could differentiate into various retinal cells types**. (E and F)** IF microscopy images of retinal cross-sections show the rescue of *yy1a* MO and *yy1b* MO effect by the transfection of respective MO binding site mutated mRNA in retina at 4dpi (E), which is quantified (F). **(G)** IF microscopy images of retinal sections at 4days post corneal injection of *yy1a* and *yy1b* mRNA, without any injury to the retina, shows no proliferating cells. Scale bars represent 10μm in (B, D, E) the asterisk marks the injury site and GCL, ganglion cell layer; INL, inner nuclear layer; ONL, outer nuclear layer in (B, D, E); dpi, days post injury. Error bars represent SD, *p < 0.0001 in (C); n.s. non-significant in (F).

**Figure S4.**
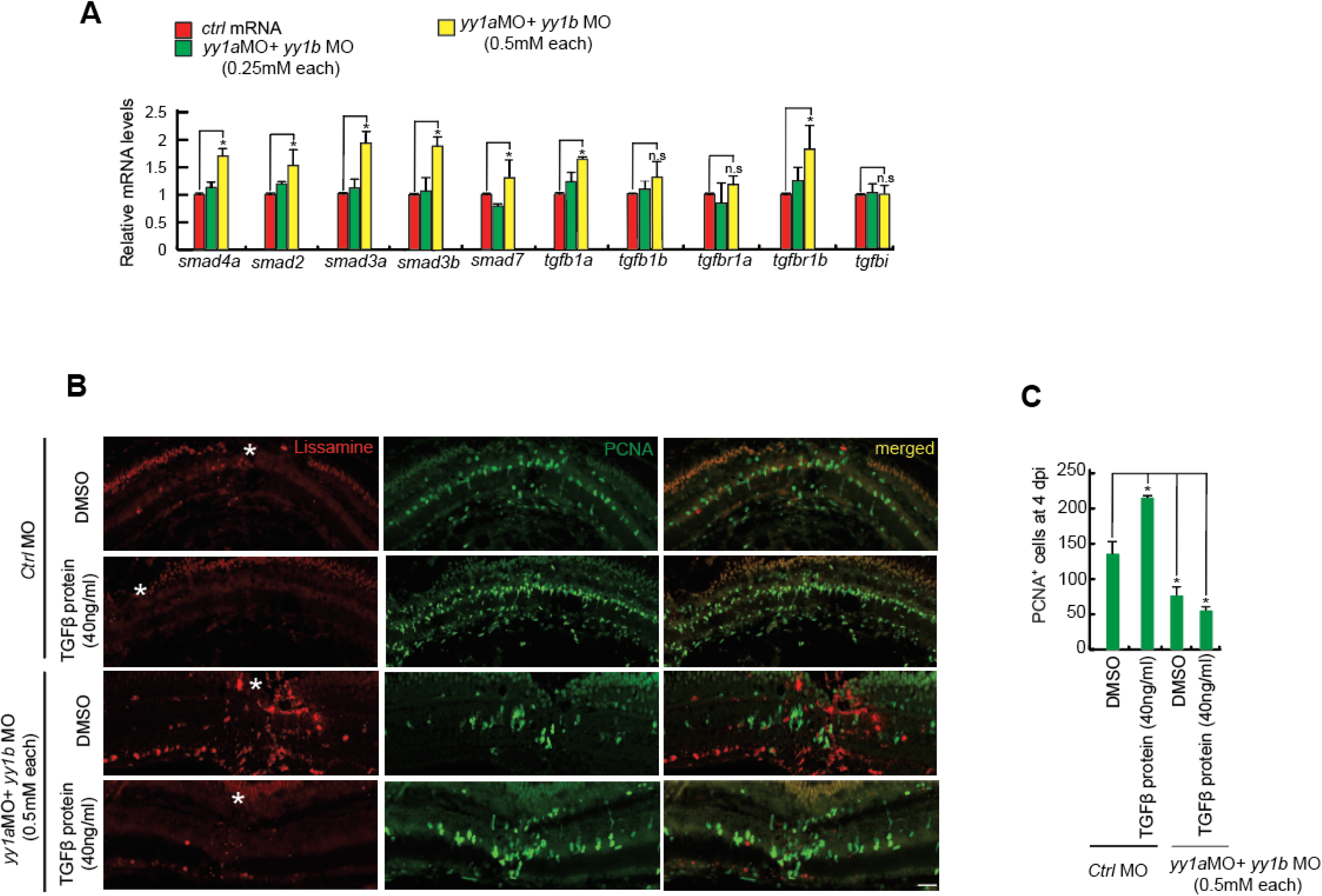
Yy1 regulates the proliferation of progenitor cells through TGFβ signalling. **(A)** qPCR analysis shows decrease in the levels of TGFβ signalling component genes in *yy1a* and *yy1b* combined knockdown condition. **(B and C)** IF microscopy images of retinal section at 4dpi, shows decline in the number of PCNA^+^ retinal progenitors in the combined treatment of *yy1a* and *yy1b* morpholino along with TGFβ protein (B) and which is quantified also (C). Error bars represent SD, *p < 0.005 in (A, C); n.s. non-significant in (A). Scale bars represent 10μm in (B).

**Figure S5.**
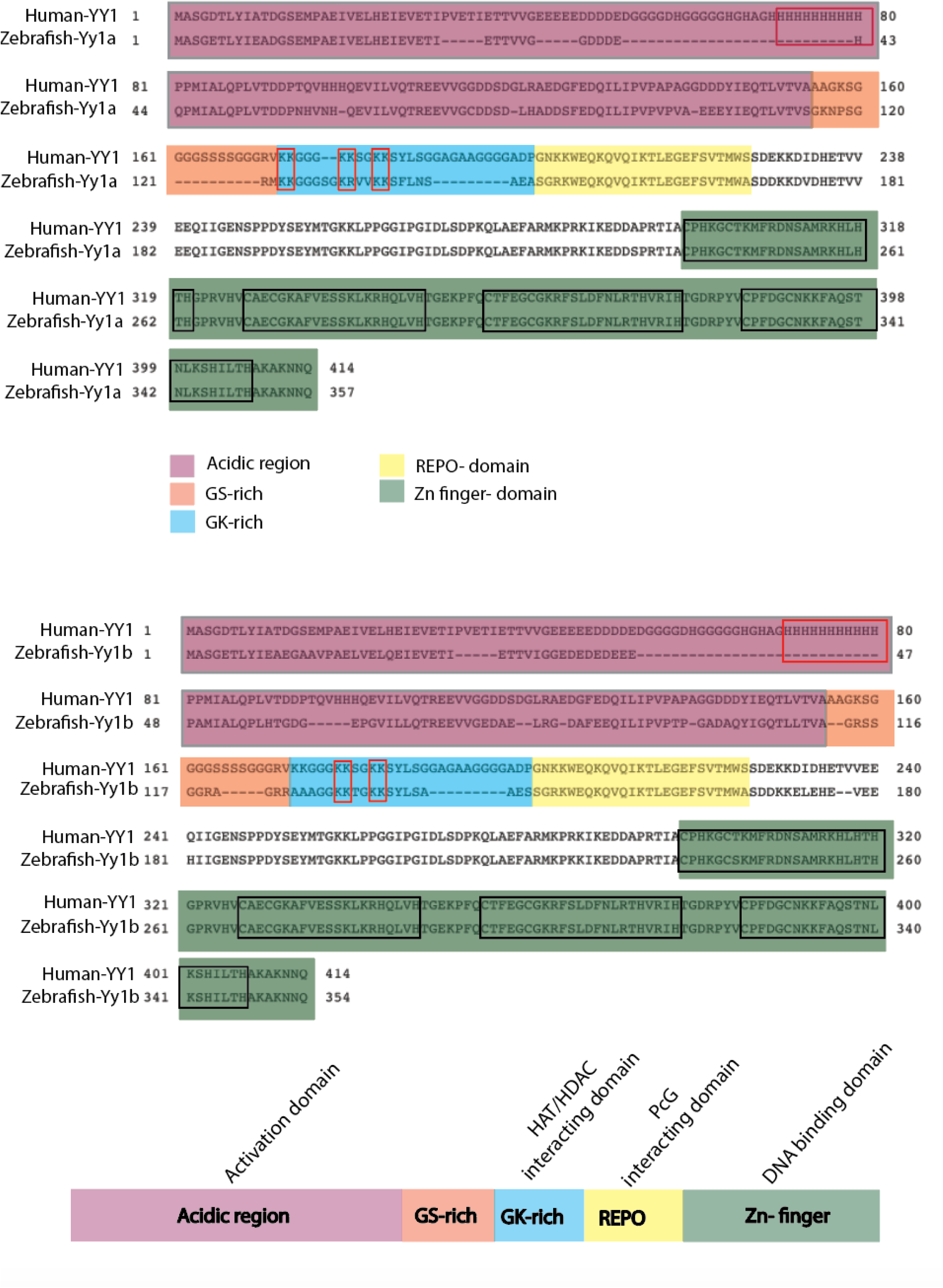
NCBI BLAST analysis of Zebrafish Yy1a and YY1b with Human YY1 shows the conserved lysine residues in the GK-rich region. **(A)** NCBI BLASTP analysis of human YY1 with Yy1a and Yy1b predicts different domains of zebrafish Yy1a (above) and Yy1b (lower). The blue region indicates the GK-rich region, which is the HAT and Hdacs interacting domain. There were six lysine residues in Yy1b while Yy1b has 4 lysine residues marked with red boxes in the blue region, which were mutated to alanine (neutral mutation) and glutamine (acetylated mimetic mutation).

**Figure S6.**
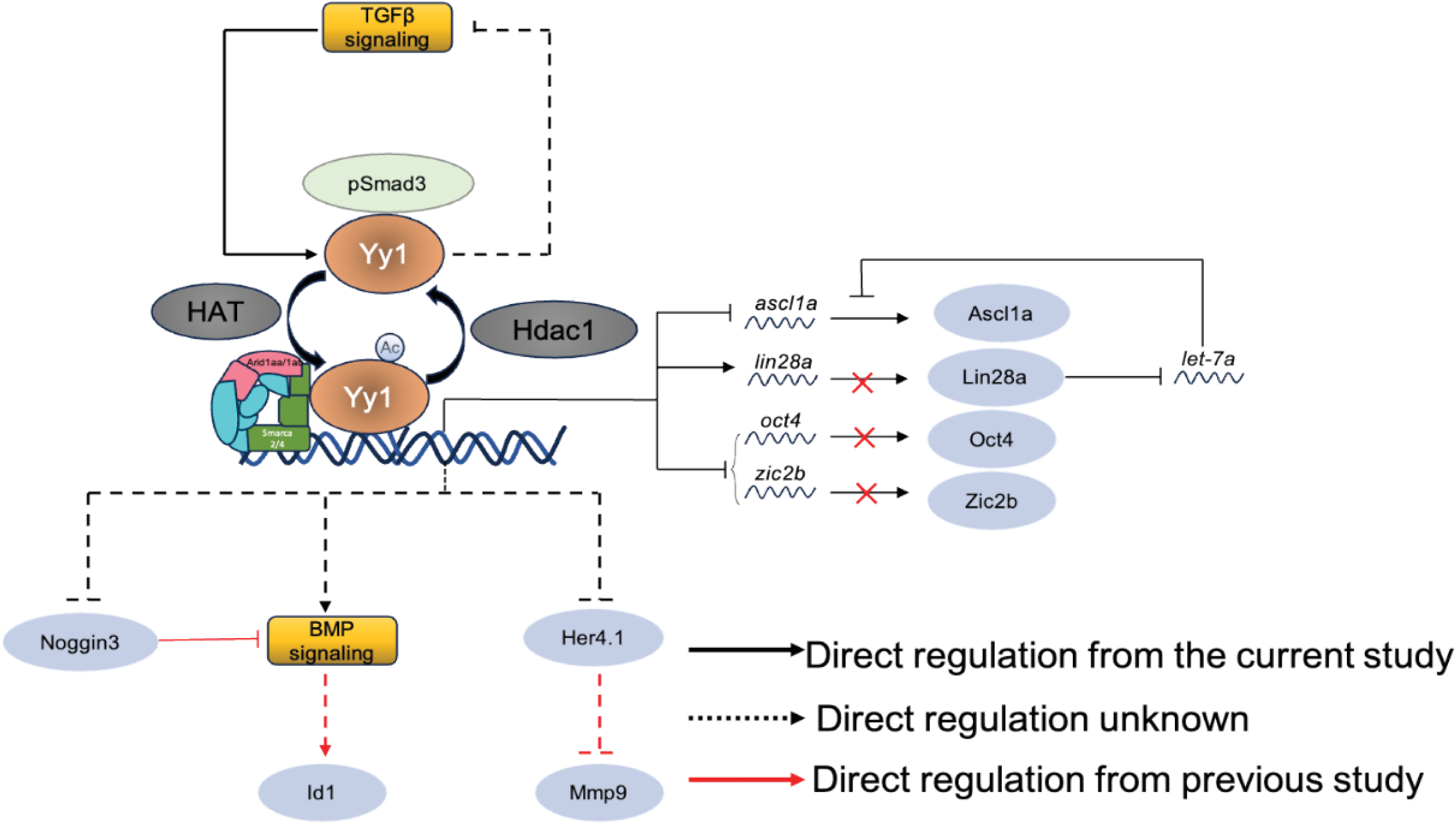
A diagrammatic model of various pathways elucidated from this study.

